# Prophase Chromosomes Relocalization to Nuclear Periphery in NPP-3/NUP205 Depletion Protects Genome Stability

**DOI:** 10.64898/2026.03.04.709728

**Authors:** Ling Jiang, Yu Chung Tse, Karen Wing Yee Yuen

**Author notes:** Correspondence (K.W.Y.Y.), (Y.C.T.).

## Abstract

The nuclear envelope (NE) mediates transport between nucleus and cytoplasm in eukaryotic cells and protects genetic materials against cytoplasmic enzymes. Nuclear pore complexes (NPCs) regulate chromosome architecture, genome integrity, gene transcription, and cell division. In *Caenorhabditis elegans* embryos, depletion of NPP-3/NUP205 causes NE rupture, premature chromosome condensation, and relocalization of condensed chromosomes to the nuclear periphery, similar to responses during anoxia and quiescence. This chromosomal relocalization depends on the spindle assembly checkpoint (SAC), inner kinetochore proteins, and partially on the NE rupture repair proteins BAF-1 and LEM-2. NPP-3 depletion prolongs prophase and prometaphase, as mediated by SAC proteins MDF-1 and MDF-2. Additionally, NPP-3 depletion alters MDF-1 localization, removing it from NE and increasing its nuclear accumulation, while reducing import of kinetochore components such as KNL-1, BUB-1, and HCP-1. In 20-30 cell-stage embryos, MDF-1 foci are observed on peripheral chromosomes during prophase. Both MDF-1 and MDF-2 accumulate on chromosomes during prometaphase. The increased incidence of lagging chromosomes, DNA damage, and micronuclei upon NPP-3 and MDF-1 depletion, suggesting that peripheral chromosome localization may serve as a protective mechanism against DNA damage. These findings shed light into cellular responses to NE rupture, with potential implications for laminopathies and cancers involving nuclear envelope defects.

## Introduction

The nuclear envelope (NE) consist of two phospholipid bilayer—the inner nuclear membrane (INM) and the outer nuclear membrane (ONM), along with associated components such as nuclear membrane proteins, the nuclear lamina, and nuclear pore complexes (NPCs) (1). The ONM is continuous with the endoplasmic reticulum (ER), and during telophase, nuclear membrane proteins and ER-derived lipid reform around chromosomes to reestablish the NE (2, 3). The INM is linked to the nuclear lamina, a meshwork of intermediate filaments primarily composed of lamin proteins that provides structural support and serves as an attachment platform for chromatin (4). Mutations in lamins lead to a spectrum of inherited disorders known as laminopathies (5, 6). The lamina is indirectly connected to the cytoskeleton through LINC (Linker of nuleoskeleton and cytoskeleton) complexes to facilitate the positioning and movement of chromosomes (7).

The nuclear envelope serves as a selectively permeable barrier that regulates molecular exchange between the nucleus and cytoplasm through specialized transport channels, most prominently the nuclear pore complexes (NPCs) (8). The NPC is positioned at the junction of the INM and ONM and can be subdivided into seven structural subcomplexes: the cytoplasmic ring, nucleoplasmic ring, cytoplasmic filament, transmembrane region, inner ring, central channel, nuclear basket and linker (9). Beyond mediating transport, the NE also participates in key cellular processes such as chromatin organization, epigenetic regulation, gene expression, DNA repair and replication, and cell cycle control (9–11).

Heterochromatin predominantly localizes to the nuclear periphery and consist of densely packed, transcriptionally silent regions enriched with modifications such as H3K9me3 and H3K27me3 (12). Its peripheral positioning contributes to gene regulation and genome stability during differentiation (13–15). The nuclear lamina and nuclear pore complexes (NPCs) are key mediators of heterochromatin anchoring, particularly inner ring and nuclear basket nucleoporins. Certain nucleoporins, known as adaptor nucleoporins, acting as bridges that connect chromatin to the NPC or serving as platforms for nuclear transport receptors and regulatory factors (16–18). For instance, the nuclear basket protein NUP153 binds across mammalian and fly genomes, as shown in multiple ChIP-seq studies, targeting silent genes, active promoters and enhancers, and CTCF-binding sites (19). Additionally, the linker component NUP93 and inner ring components NUP188 and NUP205 predominantly associate with facultative heterochromatin regions, such as promoter regions of the HOXA gene cluster, anchoring them to the nuclear periphery to promote transcriptional repression in diploid colorectal cancer (DLD1) cells (20). In *C. elegans*, depletion of the linker component NPP-13/NUP93 or inner ring component NPP-3/NUP205 disrupts transport of molecules smaller than 70 kDa, impairs cell cycle progression, and triggers abnormal chromosome condensation at the nuclear periphery (21, 22). However, the precise mechanisms by which NPP-3/NUP205 regulates chromosome localization remain unclear, and whether it specifically targets particular chromosome domains or acts globally requires further investigation of its interaction networks and functional pathways.

In *C. elegans*, which possess holocentric chromosomes lacking defined peri-centromeric regions, heterochromatin markers such as H3K9me3 are enriched along chromosome arms (23) and associate with lamina-associated domains (LADs) to promote gene repression and genome organization during differentiation (24). In the early- to mid-stage embryos, the chromodomain protein CEC-4 binds directly to H3K9me3 to anchor heterochromatin to the nuclear periphery, without affecting transcription repression (24, 25). Following differentiation, a second heterochromatin-sequestering pathway is activated. For example, MRG-1 functions as a redundant chromatin anchor in intestinal cells upon CEC-4 depletion (26).

Telomeres at chromosome ends cluster at the nuclear envelope during various cellular processes across species, including interphase in budding yeast (27), meiotic prophase in the *C. elegans* gonad, embryonic prophase (28, 29) and post-mitotic nuclear assembly in human cells (30). This clustering is mediated by the nuclear envelope protein SUN1 in humans/yeast (SUN-1 in *C. elegans*) and the telomere shelterin component RAP1 in yeast and human cells (27, 30) or telomere-binding protein POT-1 in *C. elegans* early embryos (29). During meiosis I prophase in *C. elegans*, SUN-1 localizes to the inner nuclear membrane where it interacts with pairing center proteins such as HIM-8, while the KASH domain protein ZYG-12 localizes in the outer nuclear membrane and interacts with cytoplasmic dynein (31). The SUN-1-ZYG-12 interaction transmits force generated by the cytoskeleton across the nuclear envelope to chromosomes, facilitating chromosome movement and homologous pairing (28, 32).

Structural alterations of the nuclear envelope caused by external stimuli, mechanical stress, or mutations in nuclear envelope components can trigger nuclear envelope rupture, activates repair processes, and reorganize chromatin through proteins such as Barrier-to- Autointegration Factor 1 (BAF-1) and LEM-2 (33, 34). These repair mechanisms resemble the active preassembly of the NE during mitosis (35). Similarly, in LMN-1/laminB-deficient *C. elegans* embryos, condensed chromosomes localize to the nuclear periphery, with LEM-2 accumulating at rupture sites (36).

Under respiratory stress conditions such as anoxia (oxygen deprivation) or treatment with respiratory chain inhibitors like sodium azide, nearly all chromosomes in *C. elegans* embryos relocalize to the nuclear periphery during prophase (37). Similar chromosome peripheral clustering occurs in *Drosophila* embryos (38). Notably, both organisms can fully recover from anoxia without lasting damage (37, 38), suggesting that peripheral chromosome positioning may serve as a transient protective response to respiratory stress. In *C. elegans* embryos, NPP-16 supports survival through anoxia and is associated with prophase arrest, while CDK-1 also contributes to this arrest (37). However, the mechanisms and functional significance of this chromosomal relocalization remain largely unexplored. On the other hand, the Roth lab demonstrated that the spindle assembly checkpoint is essential for anoxia-induced metaphase arrest and survival (39), though whether prophase and metaphase arrests are mechanistically linked remains unclear.

In this study, we investigated the phenotypic consequences of disrupting the inner ring component NPP-3/NUP205 in *C. elegans* embryos. NPP-3 depletion extends the duration of prophase and prometaphase. NPP-3 depletion. Notably, chromosomes in NPP-3-depleted embryos localize to the nuclear periphery, with occasional chromosome segregation defects associated with nuclear rupture. Chromosome peripheral localization in NPP-3-depleted embryos does not depend on heterochromatin or telomere anchoring pathways but instead involves centromere-kinetochore components, the spindle assembly checkpoint components, and nuclear rupture repair machinery. Spindle assembly checkpoint is necessary for the extended prophase and prometaphase observed in early embryos following NPP-3 depletion. Additionally, DNA damage checkpoints is activated in late-stage embryos after NPP-3 depletion, with increased DNA damage upon simultaneous depletion of NPP-3 and MDF-1. These results suggest that chromosome positioning at the nuclear periphery may serve a protective role, mitigating DNA damage and ensuring accurate chromosome segregation. Our findings provide new insights into the connection between nuclear envelope integrity and chromosome positioning during cell division, with implications on nuclear organization under stress conditions.

## Results

### NPP-3 depletion causes chromosomes to cluster at the nuclear periphery

To investigate the effect of nucleoporins on chromosome behavior, we depleted several representative nucleoporins from different subcomplexes: NPP-2 (cytoplasmic and nucleoplasmic ring), NPP-4 (central channel), NPP-3 and NPP-13 (inner ring), and NPP-7 (nuclear basket). Depletion of these NUPs disrupted nuclear envelope integrity, causing small but numerous ruptures along the nuclear envelope (Figure S1A). Notably, in P1prophase nuclei of two-cell stage embryos, condensed chromosomes localized closed to the nuclear periphery in *npp-3(RNAi)*, *npp-7(RNAi)* or *npp-13(RNAi)* (Figure S1A). This peripheral chromosome redistribution aligned with prior observations for depletion of the NPC inner ring components (21). In contrast, no such chromosomal localization occurred in *npp-2(RNAi)* or *npp-4(RNAi)* embryos.

To gain deeper insights into the role of inner ring subcomplex in chromosome organization, we examined the subcellular distribution of NPP-3 throughout the cell cycle. We generated a CRISPR/Cas9-engineered strain expressing mCherry-tagged NPP-3 at the endogenous locus and co-expressing GFP::H2B in live-cell imaging. In both transcriptionally quiescent P1 cells and transcriptionally active 15-cell-stage embryos, mCherry::NPP-3 predominantly localized to the nuclear envelope during interphase to prophase (Figure 1A and S1B). Upon nuclear envelope breakdown (NEBD), it adopted a characteristic two-half-circle pattern, absent in the vicinity of the centrosomes (Figure 1A and S1B)), consistent with previous observations (22). Depletion of NPP-3 by feeding RNA interference substantially reduced the mCherry::NPP-3 signal at the nuclear envelope, confirming that this CRISPR/Cas9-engineered mCherry::NPP-3 reporter faithfully reflected endogenous NPP-3 localization (Figure S1C and S1D).

**Figure 1.**
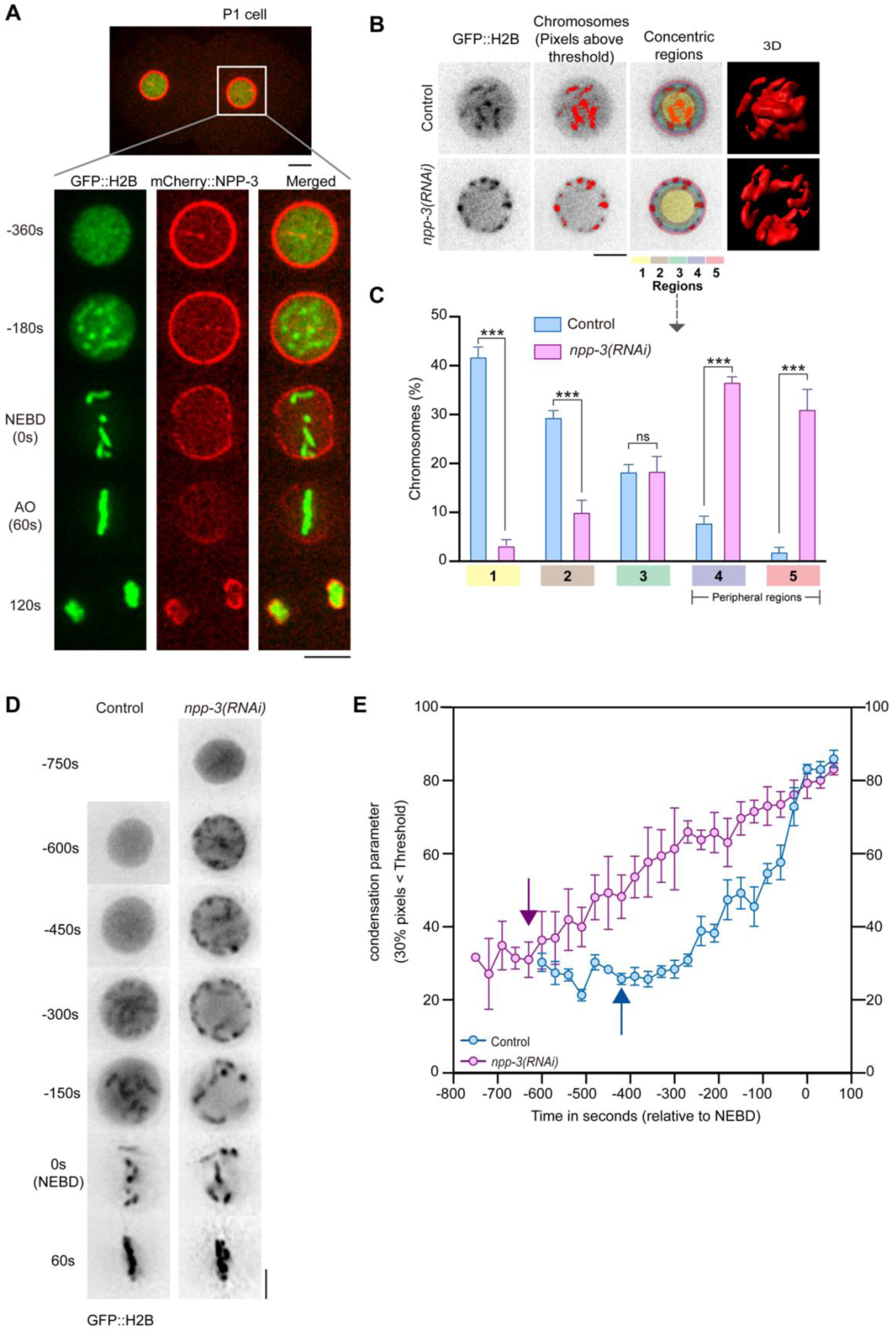
Loss of NPP-3 causes condensed chromosomes localizing to nuclear periphery. (A) Representative confocal image (of the middle single layer) of GFP::H2B and mCherry::NPP-3 in the P1 cell (white square) in two-cell stage. The P1 cell is magnified and time-lapse images from prophase to anaphase are shown. The time point 0 s represents the nuclear envelope breakdown (NEBD). AO represents anaphase onset. Scale bar, 5 μm. (B) The nucleus in the center Z plane is divided into five concentric rings or circles with equal area, as region 1-5. Regions 4 and 5 are considered as nuclear peripheral regions. A GFP::H2B intensity threshold is set to represent the chromosomes. Pixels above the threshold are assigned to Regions 1 to 5 in the control and *npp-3(RNAi)* groups. The right panel shows a snapshot of the 3-dimensional chromosome distribution after Imaris surface simulation. (C) Bar graph showing the percentage of chromosome pixels in different regions in control and *npp-3(RNAi)* groups. The sample size is 12. Error bars represent mean ± SEM. Data were analyzed using Student’s t-test; ***p<0.001; ns: not significant. (D) Representative confocal images (maximum intensity projection) of chromosome dynamics marked by GFP::H2B from prophase to metaphase in the P1 cell. The time point 0 s represents the nuclear envelope breakdown (NEBD). Scale bar, 5 μm. (E) The dynamic changes of chromosome condensation parameter, in which 30% of pixels in the ROI analyzed is below the threshold scaled intensity (<77), in the control and *npp-3(RNAi)* embryos. Blue and purple areas indicate chromosome condensation phases in control group and *npp-3(RNAi),* respectively. The sample size is 5. Error bars show mean ± SEM.

As chromosomes puncta localized to nuclear periphery after NPP-3 depletion, we hypothesize whether there is a negative regulation of NPP-3 on chromosome nuclear peripheral localization. The centrosomal kinase AIR-1/Aurora A and centrosomes regulate the loss of NPP-3 from the nuclear envelope during mitotic entry (22). Thus, NPP-3 form two-half-circles localization in prophase in wildtype. Depletion of AIR-1/Aurora A in *C. elegans* embryos results in the presence of multiple centrosomes (22, 40). The relationship between AIR-1, centrosomes and NPP-3 provides an opportunity to investigate the potential effect of AIR-1 depletion on localization of NPP-3 at the nuclear envelope. To explore whether NPP-3 localization regulates chromosomes positioning along the nuclear envelope, we depleted AIR-1/Aurora A to induce the formation of multiple centrosomes - a process associated with the disappearance of NPP-3 near the multiple centrosomes. Upon AIR-1 depletion, mCherry::NPP-3 showed fragmented localization rather than a whole circle on the nuclear envelope in the prophase of 20-30-cell stage embryos (Figure S1E). Additionally, chromosomes labeled with H2B::GFP were positioned in proximity to the nuclear envelopes, but specifically did not directly associate with the sites containing mCherry::NPP-3 foci (Figure S1E). The peaks of mCherry::NPP-3 and H2B::GFP signals exhibited an inverse relationship in *air-1(RNAi)* embryos (Figure S1F), suggesting that NPP-3 negatively regulates chromosome nuclear periphery localization. As depletion of NPP-7 also showed chromosomes localizing to nuclear periphery (Figure S1A), the relationship between NPP-7 and chromosome was also tested. Interestingly, similar to mCherry::NPP-3, chromosomes also localized at sites with low intensity of GFP::NPP-7 in *air-1(RNAi)* embryos (Figure S1G and S1H). These observations suggest that condensed chromosomes tend to localize at nuclear envelope sites without NPP-3 or NPP-7.

### NPP-3 depletion leads to earlier chromosome condensation

To uncover the underlying the aberrant chromosomal organization observed upon NPP-3 depletion, we take advantage the well-characterized mitotic dynamics of the P1 cell in *C. elegans* embryos. Live-cell imaging of GFP::H2B-labeled nuclei in *npp-3(RNAi)* embryos revealed a pronounced redistribution of all chromosomes toward the nuclear periphery (Figure 1B). This redistribution was evident in both two- and three-dimensional reconstructions generated through Imaris surface rendering (Figure 1B and 1C). To assess the functional implications of this altered chromosomal positioning induced by NPP-3 depletion, we quantified chromatin condensation dynamics in the P1 cell using the approach described by Maddox et al. (41). During prophase, a nucleus-confined square region was rescaled at each time point to normalize intensity distributions (0–255), minimizing artifacts due to changes in nuclear size and photobleaching. In control embryos, chromatin condensation began approximately 420 seconds prior to nuclear envelope breakdown (NEBD) and progressed to rod-shaped chromosomes by 120 seconds before NEBD (Figure 1D and 1E). In contrast, *npp-3(RNAi)* embryos exhibit an earlier onset of condensation, detectable as early as 630 seconds prior to NEBD, while reaching a comparable level of chromatin compaction at NEBD (Figure 1D and 1E). In terms of development, control embryos advanced to the three-fold stage within ∼15 hr after P1 division, whereas NPP-3-depleted embryos arrested during early gastrulation, with chromosomes persistently localized at the nuclear periphery (Figure S1I). This developmental arrest correlates with the embryonic lethality observed in viability assays, confirming the essential role of NPP-3 in embryogenesis (Figure S1J). Together, our findings demonstrate that NPP-3 depletion advances the timing of chromatin condensation, drives peripheral chromosome positioning, and impairs normal embryonic development.

### NPP-3 depletion results in increased heterochromatin and transcription silencing

Heterochromatin and transcriptionally repressed genes are typically positioned at the nuclear periphery. This organization is mediated by interactions between specific histone modifications and their associated binding proteins, and is characterized by the enrichment of key heterochromatic marks such as H3K9me2, H3K9me3, and H3K27me3 at the nuclear lamina (42). Given that NPP-3 depletion causes chromosome to relocalize to the nuclear periphery and triggers earlier chromatin condensation, we hypothesized that this peripheral repositioning might be associated with transcriptional repression.

To test this, we performed RNA sequencing to compare the global gene expression profiles between control and *npp-3(RNAi)* embryos. Compared to the control embryos, 30% of genes were downregulated, 2% were upregulated, and 68% remained unchanged upon NPP-3 depletion (Figure 2A), indicating large-scale transcriptional repression. Gene Ontology analysis revealed that up-regulated genes were primarily associated with mitotic cell cycle phase transitions and cyclin-dependent protein serine/threonine kinase activity (Figure S2A), whereas down-regulated genes were enriched for functions related to RNA polymerase II and larval development (Figure S2B). Notably, chromosomal mapping of both up- and down-regulated genes showed no positional bias (Figure S2C and S2D).

**Figure 2.**
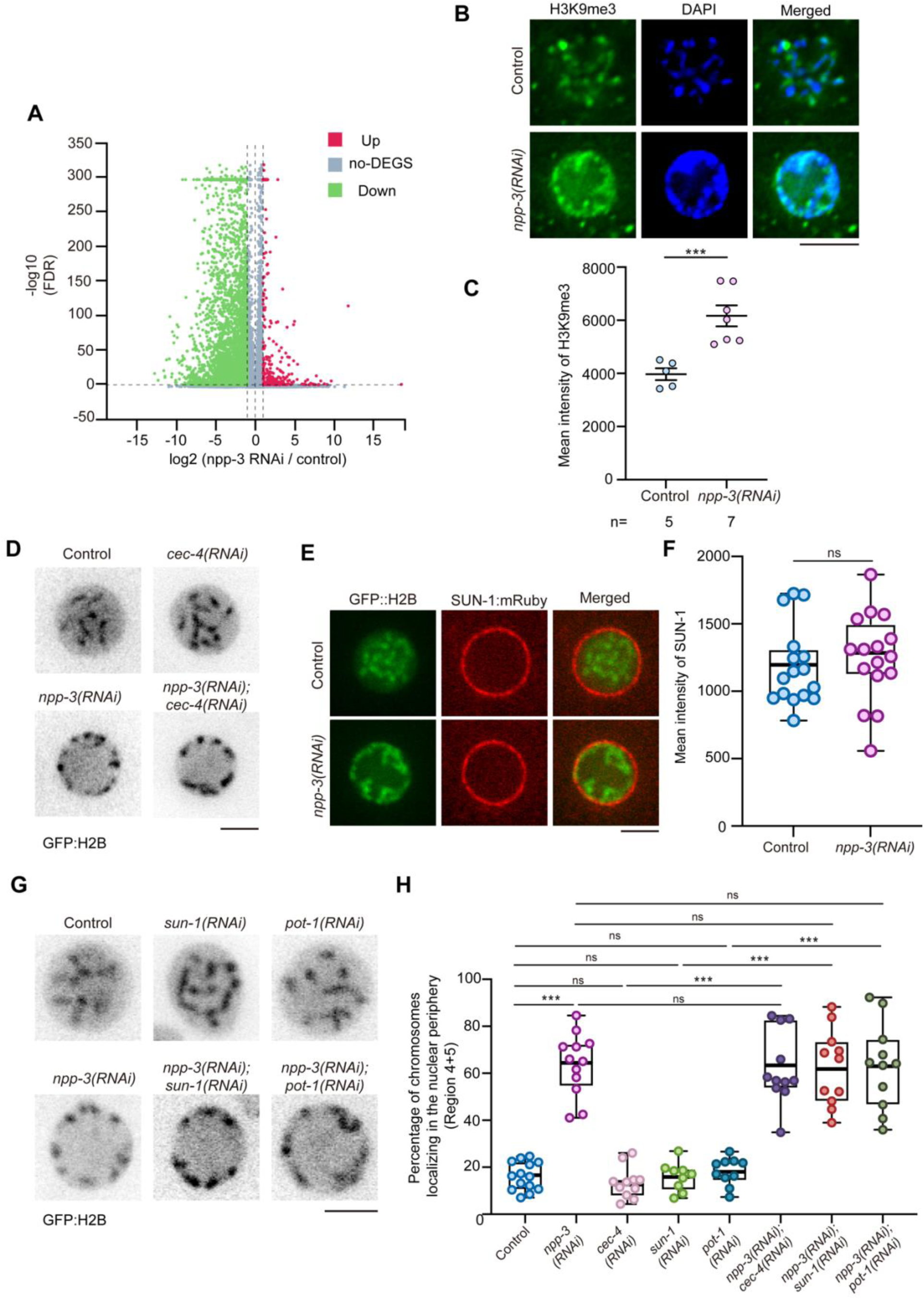
Heterochromatin and telomere anchoring pathway are not involved into peripheral chromosomal localization. (A) The volcano plot illustrates the distribution of genes based on their expression levels in the control and *npp-3(RNAi)* embryos. Up-regulated genes are shown in red, down-regulated genes in green, and genes with no significant deregulation in gray (no-DEGs). (B) Representative images of immunofluorescence staining for H3K9me3 and chromosomes (DAPI) in the P1 cell of control and *npp-3(RNAi)* embryos. Scale bar 5 μm. (C) Quantification of mean intensity of H3K9me3 in the P1 cells in (D). Each dot represents one sample. Error bars represent mean ± SEM. Data were analyzed using Student’s t-test; ***p<0.001. (D) Representative single-plane images of P1 cell with GFP:::H2B marked chromosomes in control, single and double RNAi. Scale bar 5 μm. (E) Representative images of P1 cell expressing GFP::H2B and SUN-1::mRuby in control and *npp-3(RNAi)* groups. Scale bar 5 μm. (F) Quantification of the mean intensity of SUN-1 in P1 cells of the control and *npp-3(RNAi)* embryo in the (B). Error bars represent mean ± SEM. Data were analyzed by Student’s t-test; **p<0.005. (G) Representative single plane images of P1 cell with GFP:::H2B marked chromosomes in control, single and double RNAi. Scale bar 5 μm. (H) Quantification of the chromosomal nuclear periphery localization in different conditions shown in the (D) and (G) by concentric assay shown in Figure 1B.

To further investigate the chromatin state at the nuclear periphery, we conducted immunofluorescence staining with an antibody against the heterochromatin marker H3K9me3 in fixed embryos. In control embryos, H3K9me3 foci were primarily observed on subsets of chromosomes arms/ends near the nuclear periphery. In contrast, *npp-3(RNAi)* embryos displayed a greater number of H3K9me3 foci and higher mean fluorescence intensity across most chromosomes at the nuclear periphery (Figure 2B and 2C). These findings suggest that chromosomes relocated to the nuclear periphery in NPP-3-depleted embryos adopt a more heterochromatic, transcriptionally repressed state.

### Heterochromatin and telomere anchoring pathways are not involved in peripheral chromosomal localization

In *C. elegans* embryos, the inner nuclear membrane protein CEC-4 is known to anchor H3K9-methylated heterochromatin at the nuclear periphery (43). To investigate whether CEC-4 is required for the peripheral chromosome accumulation caused by NPP-3 depletion, we examined chromosome localization following CEC-4 depletion alone and in combination with NPP-3 depletion. Depletion of CEC-4 alone did not alter chromosome positioning (Figure 2D and 2H). However, co-depletion of NPP-3 and CEC-4 resulted in a peripheral localization phenotype indistinguishable from *npp-3(RNAi)* alone (Figure 2D and 2H), demonstrating that CEC-4 is dispensable for this process.

An alternative mechanism for chromosome anchoring at the nuclear periphery involves telomere clustering mediated by the SUN-1/POT-1 pathway in *C. elegans*, which is essential for meiotic progression (29). To investigate whether the peripheral chromosome localization observed upon NPP-3 depletion depends on the SUN-1-POT-1 pathway, we analyzed SUN-1 distribution and performed genetic epistasis experiments. Our results showed that *npp-3(RNAi)* did not alter SUN-1 localization or its fluorescence intensity at the nuclear envelope (Figure 2E and 2F). Moreover, depletion of SUN-1 or POT-1 individually did not affect chromosome positioning, and their co-depletion with NPP-3 failed to alter the peripheral accumulation phenotype (Figure 2G and 2H). Collectively, these findings indicate that the peripheral chromosome localization induced by NPP-3 depletion is independent of the SUN-1/POT-1-mediated telomere anchoring pathway.

### Nuclear rupture repair is required for chromosomes localizing to nuclear periphery in NPP-3 disruption

Given that NPP-3 localizes to the nuclear envelope, we hypothesized that NPP-3 depletion might compromise nuclear envelope integrity and consequently alter chromosome status. This hypothesis is based on previous findings that nuclear envelope disruption can trigger chromosome condensation at rupture sites (33, 44) and induce DNA damage (45). To test this, we used a strain co-expressing NUP channel component NPP-1/NUP43::GFP and mCherry::H2B to visualize nuclear envelope structure and chromatin organization. Nuclear envelope permeability was further assessed using a LacI-tagged GFP reporter (46). In control embryos, the NPP-1 signal appeared continuous, indicating an intact nuclear envelope. In contrast, in *npp-3(RNAi)* embryos, the NPP-1 signal displayed numerous gaps and discontinuities (Figure 3A and 3B), signifying structural compromise of nuclear envelope. To quantify the extent of nuclear envelope damage, we performed line-scan analysis along the nuclear envelope at different embryonic stages. This analyses revealed a progressive increase in both the size and number of rupture sites from the one-cell to the multicellular stage in *npp-3(RNAi)* embryos (Figure S3C and S3D). This structural compromise was accompanied by impaired nuclear envelope barrier function, as indicated by a significant reduction in the nuclear import of the LacI-GFP reporter (Figure S3A and S3B), consistent with previous findings that NPP-3 regulates nuclear permeability (22).

**Figure 3.**
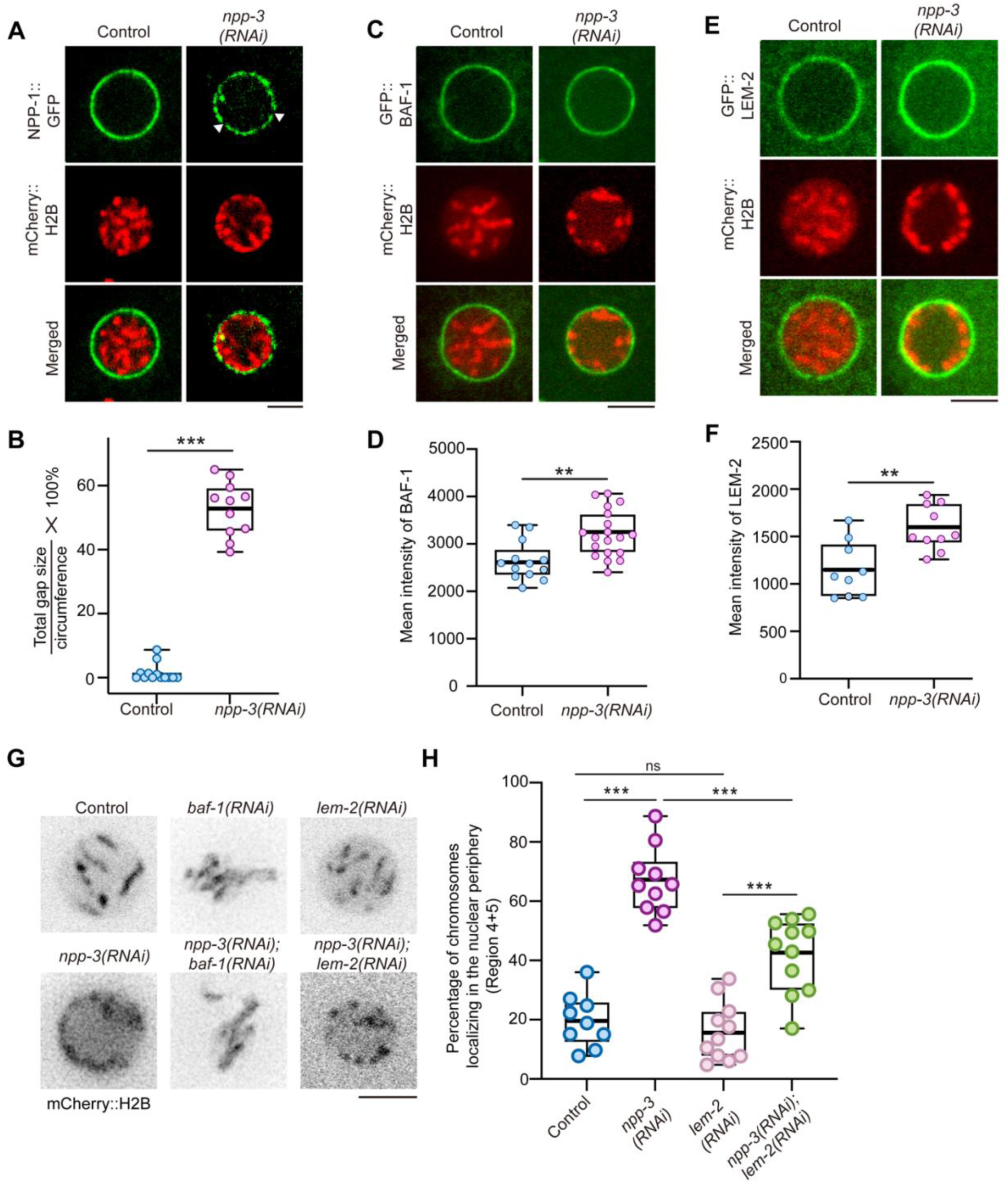
Nuclear rupture repair is required for chromosomes localizing to nuclear periphery. (A) Representative confocal images in P1 cell of the control and *npp-3(RNAi)* embryos expressing NPP-1::GFP and mCherry::H2B.. White triangles indicate the sites of nuclear envelope rupture. Scale bar 5 μm. (B) Quantification of total gap size per nucleus in the (A). Each dot represents one nucleus. Error bars represent mean ± SEM. Data were analyzed using Student’s t-test; ***p<0.001. (C) and (E) Representative confocal images in P1 cell of the control and *npp-3(RNAi)* embryos expressing GFP::BAF-1 and mCherry::H2B (C); GFP::LEM-2 and mCherry::H2B (E). Scale bar 5 μm. (D) and (F) quantification of the mean intensity of BAF-1(C) and LEM-2 (E) in P1 cells of the control and *npp-3(RNAi)* embryos. Error bars represent mean ± SEM. Data were analyzed by Student’s t-test; **p<0.005. (G) Representative images of a single plane of mCherry::H2B marked chromosomes in control, single and double RNAi. Scale bar 5 μm. (H) Quantification of the chromosomal nuclear periphery phenotype in different conditions shown in the (I) by concentric assay shown in Figure 1B.

Nuclear envelop rupture triggers a repair response involving recruitment of Barrier-to-Autointegration Factor (BAF-1) and LEM-domain protein LEM-2, both of which can promote chromatin condensation and repositioning (36, 47–49). To determine whether this repair pathway contributes to peripheral chromosome accumulation upon NPP-3 depletion, we examined the roles of BAF-1 and LEM-2. Consistent with nuclear envelope ruptures, *npp-3(RNAi)* embryos exhibited increased accumulation of both BAF-1 and LEM-2 at the nuclear periphery (Figure 3C to 3F). While *baf-1 (RNAi)* alone and *npp3(RNAi);baf-1 (RNAi)* double depletion produced severe chromosomes clustering that precluded epistatic analysis of chromosome positioning, we examined LEM-2 single and double depletion with NPP-3. *lem-2(RNAi)* alone did not alter chromosome positioning, whereas co-depletion of NPP-3 and LEM-2 partially suppressed the peripheral chromosome accumulation phenotype (Figure 3G and 3H). These findings are consistent with a model where NPP-3 depletion causes nuclear envelope rupture and BAF-1-dependent recruitment of LEM-2 to the affected regions (35), which may promote chromosome repositioning to the periphery.

Additionally, during embryonic development, nuclear rupture size increases (Figure S2D), correlating with compromised nuclear integrity and potential accumulation of DNA damage from cytoplasmic nucleases over time (50). Notably, the DNA damage sensor HUS-1 (a 9-1-1 complex component) colocalized with chromosomes in *npp3(RNAi)* in 20 to 30-cell-stage embryos but not in P1 cell (Figure S3E and S3F). These HUS-1 foci indicated activation of the DNA damage checkpoint during 20-30 cell stage embryogenesis upon NPP-3 depletion. Additionally, double depletion of NPP-3 and CHK-1 did not revert NPP-3-induced peripheral chromosome localization in P1 cells (Figures S3G and S3H), confirming that this relocalization is independent of the DNA damage checkpoint pathway.

### Spindle assembly checkpoint and centromere-kinetochore proteins are required for chromosomes localizing to nuclear periphery in NPP-3 disruption

Under respiratory stress conditions such as anoxia or sodium azide treatment, *C. elegans* embryos exhibit a distinctive prophase phenotype in which nearly all chromosomes localize near the nuclear periphery (37). Loss of nucleoporin NPP-16/NUP50 suppresses prophase arrest under anoxia, suggesting a role for nucleoporins in this process. The nuclear envelope localization of the spindle assembly checkpoint (SAC) component MDF-1/MAD1 depends on interactions with nuclear pore complex NUP153 in human cells (51) and NPP-5/NUP107 in *C. elegans* (52). Such nuclear envelope localization of MAD1 has been shown to contribute to SAC activation before nuclear envelope breakdown (53, 54).

On the other hand, the SAC plays a crucial role in anoxia-induced metaphase arrest and survival of *C. elegans* embryos (39). Oxygen deprivation also induces chromosome relocalization toward the nuclear periphery (38, 55). Moreover, in *Drosophila* embryos, SAC components such as MPS1 and BUB3 accumulate on chromosomes at the spindle midzone during metaphase and subsequently relocalize to the centrosomes during interphase under anoxic conditions (56). Although the SAC does not appear to be required for anoxia-induced prophase arrest (39), it remains unclear whether the SAC can function during interphase and prophase, and if it is involved in controlling chromosome positioning following anoxia treatment.

To investigate whether SAC components and centromere-kinetochore contribute to the peripheral chromosome localization observed upon NPP-3 depletion, we performed epistatic analysis of chromosome localization. Individual depletions were carried out for SAC proteins MDF-1/MAD1, MDF-2/MAD2, SAN-1/MAD3, and for inner kinetochore proteins HCP-3/CENP-A, HCP-4/CENP-C, and KNL-1, as well as outer kinetochore proteins NDC-80, BUB-1, BUB-3, and HCP-1/CENP-F, both alone and in combination with *npp-3(RNAi)*. Chromosomal positioning during prophase in P1 cells was assessed. Efficient depletion of *npp-3* RNA was verified by NPP-3::mCherry signal (Figure S4A and S4B). We found that depletion of MDF-1 or MDF-2 alone did not affect chromosomal positioning; however, co-depletion with NPP-3 abolished peripheral chromosome localization, restoring central nuclear localization comparable to controls (Figures 4A and 4B). On the other hand, depletion of inner kinetochore components (HCP-3, HCP-4, KNL-1) did not alter prophase chromosome positioning, although it disrupted chromosome segregation (Figures 4A and S4C). Notably, co-depletion of HCP-3 or HCP-4 with NPP-3 restored central chromosome positioning, whereas co-depletion of NPP-3 and KNL-1 retained partial peripheral chromosomal localization, suggesting distinct contributions of these kinetochore components to chromosome–nuclear envelope interactions..

**Figure 4.**
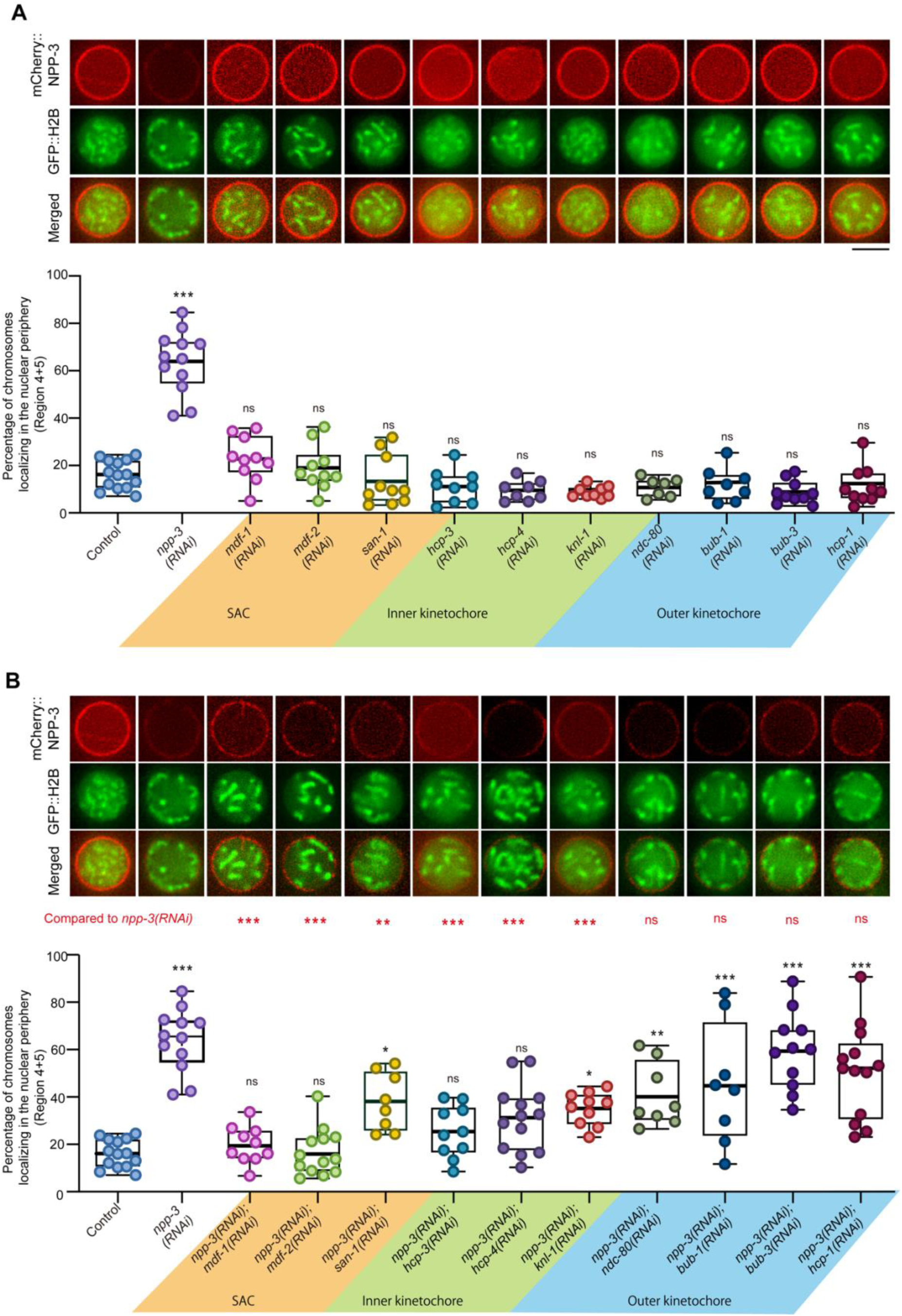
Spindle assembly checkpoint, centromere and kinetochore proteins are required for chromosomes localizing to nuclear periphery in NPP-3 disruption. (A) The upper panel is the representative images of a single plane of P1 cell expressing GFP::H2B and mCherry::NPP-3 in control, different single RNAi. Scale bar 5 μm. The bottom panel is the quantification of the chromosomal nuclear periphery phenotype in different conditions shown in the upper panel by concentric assay shown in Figure 1B. (B) The upper panel is the representative images of a single plane of P1 cell expressing GFP::H2B and mCherry::NPP-3 in control, different double RNAi corresponding to (A). Scale bar 5μm. The bottom panel is the quantification of the chromosomal nuclear periphery phenotype in different conditions shown in the upper layer by concentric assay shown in Fig.1B. Data were analyzed by one-way ANOVA (and nonparametric or mixed); *p<0.05, ***p<0.001, ns: not significant when compared with control. Upper lane marked by red comparing with *npp-3(RNAi)* group. Bottom lane is comparing with control group.

Conversely, depletion of outer kinetochore proteins (NDC-80, BUB-1, BUB-3, HCP-1) did not alter prophase chromosome positioning in P1 cells, although loss of NDC-80 or BUB-1 caused embryonic lethality at later developmental stages (57, 58). Importantly, co-depletion of these outer kinetochore proteins with NPP-3 did not alter the peripheral chromosome localization phenotype. Collectively, these findings reveal that SAC proteins MDF-1, MDF-2 and SAN-1, together with inner kinetochore components, play essential roles in regulating chromosome positioning at the nuclear periphery during prophase.

### Absence of NPP-3 Activates the Spindle Assembly Checkpoint to extend both prophase and prometaphase

As SAC and kinetochore proteins are required for the peripheral chromosomal localization upon NPP-3 depletion, we wonder whether SAC is activated in prophase and prometaphase. Firstly, we monitored nuclear envelope and chromatin dynamics in the P1 cell by using a strain expressing BAF-1::GFP; mCherry::H2B. BAF-1 marks the nuclear membrane and serves as an indicator of NE remodeling timing (59), while also highlighting anaphase-bridged chromatin in *lmn-1(RNAi)* embryos (60). We found that NPP-3 depletion causes a significant delay in the cell cycle. Specifically, the duration of prophase—measured from nuclear envelope reassembly (NER) in the P0 cell to nuclear envelope breakdown (NEBD) in P1 cell—was prolonged to 950.0 ± 112.1 seconds, compared with 782.4 ± 75.1 seconds in control embryos (Figure 6A and 6B). Additionally, the interval between NEBD and anaphase onset (AO) was 156.0 ± 25.0 seconds in *npp-3(RNAi)* embryos, more than doubled of the duration in controls (79.5 ± 14.7 seconds) (Figure 5A and 5C).

**Figure 5.**
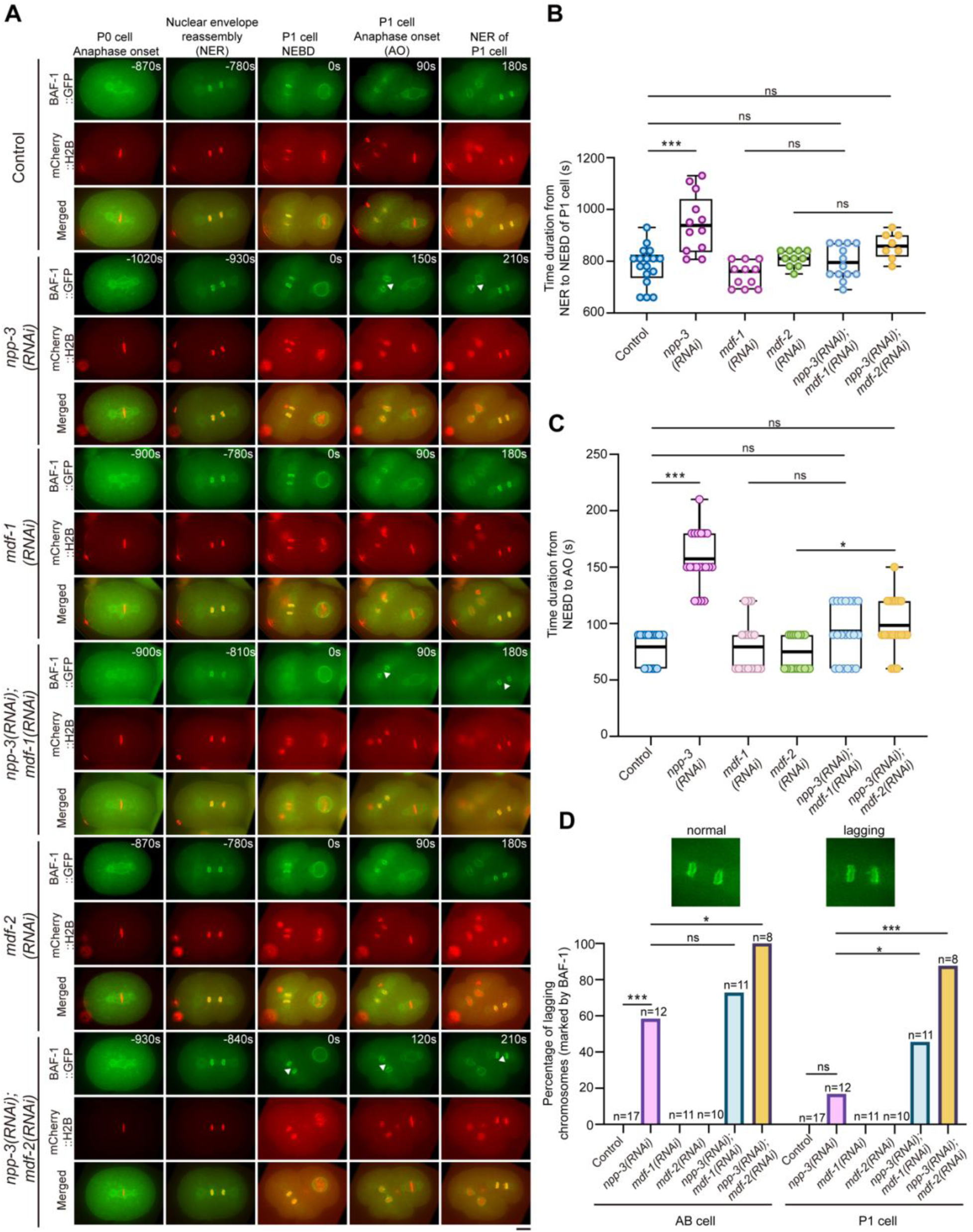
Absence of NPP-3 causes extended prophase and metaphase through MDF-1 and MDF-2. (A) Representative single-plane confocal image of P1 cell division from anaphase pf P0 cell to anaphase in control, *npp-3(RNAi)*, *mdf-1(RNAi), mdf-2(RNAi), npp-3(RNAi);mdf-1(RNAi)* and *npp-3(RNAi);mdf-2(RNAi)* embryos BAF-1::GFP and mCherry::H2B. Triangles indicate the lagging chromosomes marked by lagging BAF-1. Scale bar 10 μm. (B) Quantification of the prophase duration of P1 cell from P0 cell nuclear envelope reassembly (NER) to P1 cell nuclear envelope breakdown(NEBD) in (A). Each dot represents one embryo. Error bars represent mean ± SEM. Data were analyzed by one-way ANOVA (and nonparametric or mixed); *p<0.05, ***p<0.001, ns: not significant. (C) Quantification of the duration of P1 cell from nuclear envelope breakdown (NEBD) to anaphase onset (AO) in (A). Each dot represents one embryo. Error bars represent mean ± SEM. Data were analyzed by one-way ANOVA (and nonparametric or mixed); *p<0.05, ***p<0.001, ns: not significant. (D) Percentage of lagging chromosomes marked by BAF-1 in different conditions shown in (A). Data were analyzed by Fisher’s test; *p<0.05, ***p<0.001, ns: not significant.

**Figure 6.**
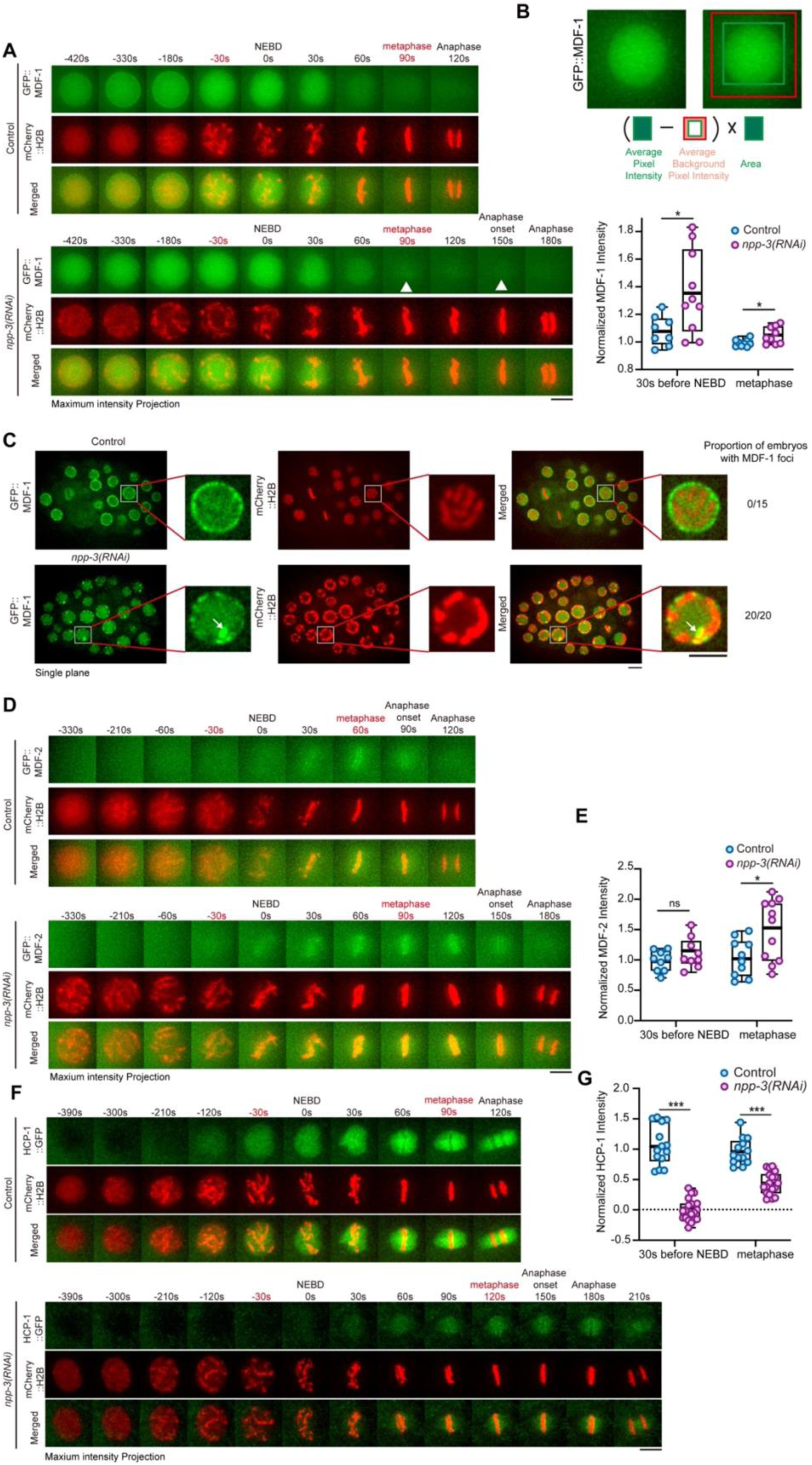
NPP-3 depletion disrupts early kinetochore assembly by reduction the import of HCP-1. (A) Representative time-lapse image (maximum intensity projection) of GFP::MDF-1 and mCherry::H2B in the P1 of control and *npp-3(RNAi)* embryos. White triangles indicate MDF-1 accumulation on chromosomes. Scale bar 5 μm. (B) Quantification of MDF-1 integrated intensity at 30 s before NEBD and metaphase in (A). In the prolonged metaphase upon NPP-3 depletion, “metaphase” is defined as the timepoint in which the highest intensity of MDF-1 is observed. Error bars represent mean ± SEM. Data were analyzed using student’s t-test, *p<0.05. (C) Representative confocal images of (a single plane of) later embryos (>25 cells) expressing GFP::MDF-1 and mCherry::H2B in control and *npp-3(RNAi)* group. White arrow indicates the MDF-1 foci. Scale bar 5 μm. (D) Representative time-lapse image (maximum intensity projection) of GFP::MDF-2 and mCherry::H2B in the P1 of control and *npp-3(RNAi)* embryos. Scale bar 5 μm. (E) Quantification of MDF-2 integrated intensity at metaphase. Error bars represent mean ± SEM. Data were analyzed using student’s t-test, *p<0.05. (F) Representative time-lapse image (maximum intensity projection) of HCP-1::GFP and mCherry::H2B in the P1 of control and *npp-3(RNAi)* embryos. Scale bar 5 μm. (G) Quantification of HCP-1 integrated intensity at 30s before NEBD and metaphase. Error bars represent mean ± SEM. Data were analyzed using student’s t-test, *p<0.05.

The prolonged NEBD-to-AO interval is a characteristic hallmark of spindle assembly checkpoint (SAC) activation (61). To determine whether this delay is SAC-dependent, we co-depleted NPP-3 with a key SAC component, MDF-1/MAD1 or MDF-2/MAD2. Single depletion of MDF-1 (79.4 ± 23.6 s) or MDF-2 (72.0 ± 15.4 s) alone did not affect the NEBD-to-AO timing compared to controls (79.5 ± 14.7 s) (Figure 5A and 5C). However, co-depletion of NPP-3, alongside MDF-1 or MDF-2, significantly reversed the anaphase onset delay, reducing the interval to 91.5 ± 24.8 seconds for *npp-3;mdf-1(RNAi)* and 97.5 ± 23.6 seconds for *npp-3;mdf-2(RNAi)* (Figure 5A and 5C). Similarly, double depletion of NPP-3 with KNL-1 also abolished the delay, whereas depletion of other kinetochore or spindle assembly proteins, including SAN-1, HCP-3, HCP-4, NDC-80, BUB-1, BUB-3, and HCP-1, did not (Figure S4D). These findings indicate that the late mitotic delay caused by NPP-3 loss is mediated through SAC activation involving MDF-1, MDF-2, and KNL-1. Notably, although bypassing the SAC restores cell cycle timing, it exacerbates genomic instability. Co-depletion of NPP-3 with either MDF-1 or MDF-2 significantly increased the frequency of lagging chromosomes, as indicated by lagging BAF-1, compared to depletion of NPP-3 alone (Figure 5A and 5D).

Interestingly, co-depletion of NPP-3 and MDF-1 also abolished the prolonged prophase duration (793.8 ± 64.4 s) caused NPP-3 depletion (950.0 ± 112.1s) to levels comparable with controls (782.4 ± 75.1s), whereas co-depletion with MDF-2 resulted in only partial restoration (832.5 ± 66.5 s) (Figure 5C). Here, we show that the SAC components MDF-1 and MDF-2 have a role in prolonging prophase in NPP-3 depletion, which is a novel role beyond their known functions in prometaphase extension.

### NPP-3 depletion causes the redistribution of SAC and kinetochore proteins during prophase

To investigate whether NPP-3 depletion activates SAC by promoting the recruitment of MDF-1/MAD1 and MDF-2/MAD2 to chromosomes from prometaphase to metaphase (58, 62, 63), We examined the dynamic localization of GFP-tagged MDF-1 and MDF-2 in control and *npp-3(RNAi)* embryos expressing mCherry::H2B. In control embryos, MDF-1 localized primarily to the nucleoplasm with mild nuclear envelope association during prophase; its signal diminished after NEBD and became undetectable on chromosomes by metaphase (Figure 6A). In contrast, NPP-3 depletion abolished MDF-1 nuclear envelope localization during prophase and caused moderate chromosome accumulation between NEBD and metaphase. Nuclear MDF-1 intensity in *npp-3(RNAi)* P1 cells showed a modest increase at ∼30 s before NEBD and at metaphase, when normalized to controls as described previously (64) (Figure 6A and 6B).

Compared to previous findings where MDF-1 aligns on chromosomes during monopolar spindles in *zyg-1(RNAi)* P1 cells, defective in centrosome duplication (62), we observed only a slight increase in MDF-1 intensity and no clear accumulation at metaphase in *npp-3(RNAi)* P1 cells. Potentially, SAC activity is weaker in the P1 cell upon NPP-3 depletion, as compared to ZYG-1 depletion with completely unattached kinetochores. Since SAC activity increases during embryogenesis with rising nuclear-to-cytoplasm ratio (65), we investigated whether MDF-1 exhibited any notable changes during embryonic development. MDF-1 intensity was also markedly elevated before NEBD and at metaphase in the EMS cell after NPP-3 depletion compared to controls, with bright MDF-1 foci co-localizing with chromosomes during prometaphase (Figure S5A and S5B). Furthermore, in the prophase of 25-cell stage, MDF-1 showed clear NE localization with diffuse signal inside the nucleus in the control embryos. Notably, bright MDF-1 foci appeared on chromosomes during prophase in embryos beyond the 25-cell stage upon NPP-3 depletion, contrasting to the nuclear envelope localization in controls (Figure 6C). These results demonstrate that NPP-3 depletion disrupts MDF-1 localization dynamics from prophase through metaphase, suggesting a potential additional role for MDF-1 during prophase.

MDF-2, another key component of spindle assembly checkpoint, exhibited a weak signal at NEBD followed by progressive chromosomal accumulation that peaked at metaphase in the P1 cell of control embryos. While there is no difference in MDF-2 signal in *npp-3(RNAi)* and control embryos at 30 seconds before NEBD in P1 cells, MDF-2 signal was slightly enhanced in the *npp-3(RNAi)* embryos compared to controls at metaphase (Figure 6D and 6E). Similar patterns were observed in EMS cells in *npp-3(RNAi)*, which showed more pronounced metaphase chromosomal accumulation (Figure S5C and S5D).

Additionally, the localization pattern of the spindle assembly checkpoint protein SAN-1 is also altered. In control P1 cells, SAN-1 entered the nucleus after NEBD, whereas upon NPP-3 depletion, SAN-1 translocated into the nucleus prior to NEBD (Figures S5E and S5F). Importantly, SAN-1 did not accumulate on chromosomes in either control or *npp-3(RNAi)* groups, consistent with previous findings that SAN-1 is not recruited to chromosomes even upon SAC activation (58).

Collectively, these data suggest that NPP-3 depletion promotes the recruitment of MDF-1 and MDF-2 to chromosomes, potentially leading to spindle assembly checkpoint activation.

To investigate whether NPP-3 depletion impacts kinetochore assembly in *C. elegans*, we examined the dynamics of inner and outer kinetochore proteins—HCP-3, KNL-1, BUB-1, and HCP-1—from prophase through anaphase. HCP-3 fluorescence intensity at 30 seconds prior to NEBD and during metaphase was comparable between control and *npp-3(RNAi)* embryos (Figure S5G and S5H). In contrast, levels of KNL-1, BUB-1, and HCP-1 were significantly reduced at 30 seconds before NEBD in NPP-3-depleted embryos (Figures S5I to S5L and 6G). At metaphase, KNL-1 accumulated on chromosomes to similar intensities in both conditions (Figure S5J), whereas BUB-1 showed increased chromosomal association in *npp-3(RNAi)* embryos compared to control (Figure S5L), consistent with SAC activation via BUB-1 recruitment (66). HCP-1 levels, however, remained diminished at metaphase under NPP-3 depletion compared to control (Figure 6F and 6G), suggesting incomplete kinetochore assembly. Collectively, these results suggest that NPP-3 may be critical for the import of kinetochore components during prophase, and its depletion hampers kinetochore assembly but activates SAC during prophase and prometaphase.

### Chromosome Anchoring at the Nuclear Periphery and Spindle assembly checkpoint MDF-1 are crucial for maintaining chromosome stability in NPP-3 depletion

Loss of the SAC abrogates the peripheral chromosomal localization induced by NPP-3 depletion during prophase (Figure 4A and 4B) and the prophase and prometaphase extension (Figure 5A and 5C). However, loss of the SAC also exacerbates genomic instability in NPP-3 depletion. In *npp-3* RNAi, even with the SAC activation, lagging chromatin was observed in the AB (58.33%) and P1 (16.67%) cells treated compared to 0% in control group (Figure 5D). Co-depletion of NPP-3 with either MDF-1 or MDF-2 increased the frequency of lagging chromosomes compared to NPP-3 depletion alone in P1 cells, increasing to 45.45% for *npp-3;mdf-1(RNAi)* and 87.50% for *npp-3;mdf-2(RNAi)* (Figure 5A and 5D). Taken together, these findings indicate that SAC activation in response to defects caused by NPP-3 loss serves to prevent premature anaphase onset, with SAC components playing specific roles in regulating mitotic phases.

To investigate whether this peripheral chromosome positioning and extended cell cycle in NPP-3 depletion serves a protective function, we compared the effects of NPP-3 depletion alone versus the double depletion NPP-3 and MDF-1 on the chromosomes. We quantified the micronuclei, marked by GFP::BAF-1, and DNA damage, labelled by HUS-1::GFP (67, 68) in 20-30 cell-stage embryos. Double depletion of NPP-3 and MDF-1 resulted in micronuclei formation in 50% of embryos, a phenotype that was absent in single depletion conditions (Figures S6A and 7A). Following NPP-3 depletion, an average of 2.40 ± 1.62 HUS-1::GFP foci co-localized with chromatin at the 20–30 cell stage, compared with 0.045 ± 0.21 and 0.075 ± 0.27 in control and *mdf-1(RNAi)* groups, respectively (Figures S6B, 7B and 7C). Double depletion of NPP-3 and MDF-1 further increased HUS-1 foci to 3.34 ± 2.39 (Figures 7B and 7C).

**Figure 7.**
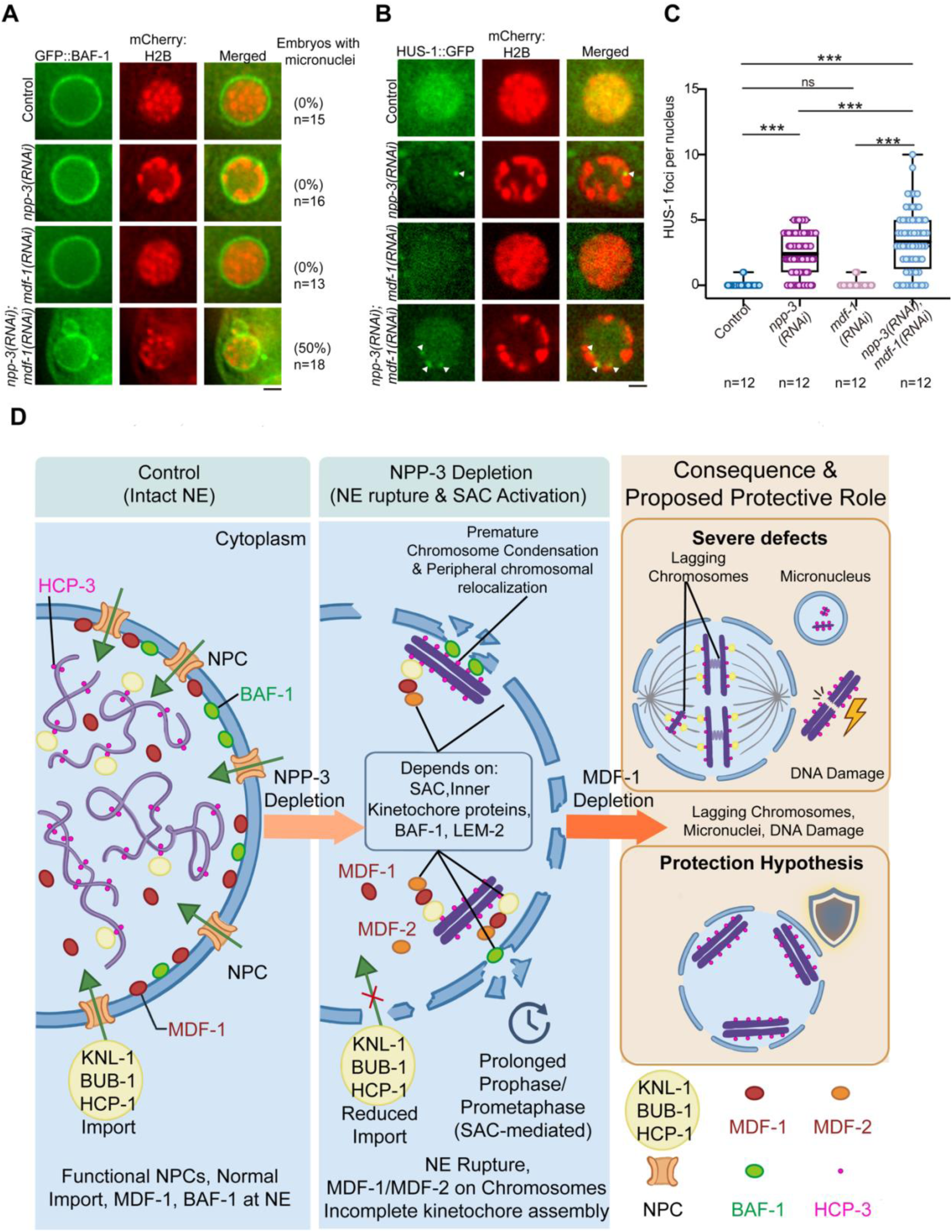
Loss of MDF-1 in NPP-3 depletion leads to more nuclear fragmentation and DNA damage. (A) Representative single-plane confocal images of nuclei from late-stage embryos (about 20-30 cell stage). expressing GFP::BAF-1 and mCherry::H2B in control, *mdf-1(RNAi)*, *npp-3(RNAi)*, and *npp-3(RNAi)*; *mdf-1(RNAi)* conditions. Scale bar 2 μm. (B) Representative images of HUS-1 and chromosome localization in the embryonic stage (about 20-30 cell stage). Arrowhead indicates the colocalization of HUS-1 foci and chromosome. Scale bar 2 μm. (C) Quantification of HUS-1 foci per nucleus including all nuclei in the embryos in different conditions in (B). Error bars represent mean ± SEM. Data were analyzed by one-way ANOVA (and nonparametric or mixed); *p<0.05, ***p<0.001, ns: not significant. (D) During mid-prophase, NPP-3 depletion causes chromosomes to localize to the nuclear periphery, depending on some spindle assembly checkpoint (SAC) components (MDF-1, MDF-2), inner kinetochore proteins, BAF-1, and LEM-2. This response prevents lagging chromosomes, DNA damage and micronuclei formation. SAC components MDF-1 and MDF-2 are also required for the prophase extension observed in NPP-3 depletion. During prometaphase, import of kinetochore proteins KNL-1, BUB-1 and HCP-1 is reduced and more MDF-1, MDF-2 and BUB-1 accumulate on chromosomes. The SAC is activated, resulting in an extended interval from nuclear envelope breakdown (NEBD) to anaphase onset when NPP-3 is depleted.

Consistent with these observations, embryonic viability was markedly reduced in the *npp-3(RNAi)* group (5.71% ± 2.48) relative to control (99.55% ± 0.78), whereas MDF-1 depletion alone had minimal effect (98.60% ± 1.95) (Figure S6C). Importantly, combined depletion of NPP-3 and MDF-1 produced a further significant decline in viability (1.38% ± 0.75), supporting the conclusion that loss of peripheral chromosome anchoring exacerbates genomic instability and severely impairs embryonic survival (Figure 7D).

## Discussion

Here, we describe the phenotype of global chromosome localization to the nuclear periphery caused by NPP-3 depletion in *C. elegans* embryos. Our candidate gene screening results indicate that this localization depends on the spindle assembly checkpoint proteins, centromere and kinetochore proteins, and nuclear rupture repair factors, but not the heterochromatin or telomere anchoring pathways. Additionally, NPP-3 depletion impairs kinetochore assembly in prophase, extends prophase and prometaphase depending on the spindle assembly checkpoint (SAC) in embryos, and triggers a DNA damage response in late-stage embryos. Notably, we observed MDF-1 and HUS-1 foci on some chromosomes during prophase. In anoxia, chromosomes also localize to the nuclear periphery, and this arrest is reversible (37). Such chromosome relocalization response in anoxia could potentially reduce transcriptional activity, as in NPP-3 depletion, and lower energy consumption. We propose that the peripheral chromosome localization in NPP-3 depletion is not a mere defect but a protective mechanism to maintain genome stability (Figure 7D).

### Not all nuclear rupture induces chromosomes localizing to the nuclear periphery

The nuclear envelope (NE) comprises several essential components, including the nuclear pore complex (NPC), nuclear lamina, and associated membrane proteins. Mutations in nucleoporins or lamins have been linked to nuclear rupture phenotypes that contribute to developmental abnormalities and a range of diseases. For example, Emery-Dreyfuss muscular dystrophy (EDMD) is associated with nucleoporin mutations (69), while laminopathies are resulted from lamin gene mutations (70). Additionally, disruptions in NE components have been implicated in neurodegenerative disorders such as Huntington’s disease and Alzheimer’s disease (18), highlighting the critical role of nuclear envelope integrity in cellular function and organismal development nuclear integrity.

In *C. elegans* embryos, lamin LMN-1 deficiency leads to prominent nuclear ruptures, accumulation of the inner nuclear membrane protein LEM-2 at rupture sites, and subsequent chromosome condensation at rupture sites, which results in the redistribution of all chromosomes to the nuclear periphery (36), similar to what we observed in NPP-3 depletion.

The depletion of the selected NUPs led to disruptions in the nuclear envelope integrity, as we observed small yet numerous nuclear rupture sites along the nuclear envelope (Figure S1A). Surprisingly, we further found that, at prophase, the condensed chromosomes were localized closed to the nuclear periphery in *npp-3(RNAi)*, *npp-13(RNAi)* or *npp-7(RNAi)* in two-cell embryos (Figure S1A). However, such chromosomal localization was not observed in *npp-2(RNAi)* or *npp-4(RNAi).* As *npp-3(RNAi)*, *npp-4(RNAi)* or *npp-13(RNAi)* embryos share similar permeability defects according to the dynamic of LacI (Figure S7), these results reveal an intriguing observation that chromosomal localization at the nuclear periphery is not directly correlated with the extent of permeability defect caused by NUPs depletions, but more specifically related to disruption of certain NPC subcomplexes in the inner ring and nuclear basket.

### Potential relationship between BAF-1 and centromere-kinetochore proteins

According to the screening, nuclear rupture repair (BAF-1, LEM-2) and centromere-kinetochore proteins (HCP-3/CENP-A, HCP-4/CENP-C, KNL-1) are involved in nuclear peripheral chromosome localization caused by NPP-3 depletion. we propose that BAF-1 may serve as a crucial molecular link between the nuclear envelope and centromere-kinetochore complexes during chromosome localization to nuclear periphery, particularly under the condition of NPP-3 depletion. Barrier-to-Autointegration Factor (BAF) is a conserved nuclear envelope (NE) component that binds chromatin and helps chromosome anchoring to the NE (47). Additionally, BAF-1 was reported to interact with centromeric protein CENP-C, for proper kinetochore assembly in metaphase which in turn regulate the chromosome segregation accuracy in *Drosophila* (44, 71). The observed decrease in CENP-C and CENP-A levels upon BAF-1 depletion supports this notion, indicating that BAF-1 might directly or indirectly influence the recruitment or maintenance of these centromeric proteins during in metaphase (71). Therefore, the potential relationship between BAF-1 and centromere-kinetochore proteins may be integral to maintaining chromosomal stability, especially when nuclear envelope integrity is compromised. Further experimental validation is warranted to elucidate the mechanistic details of this interaction during cell division.

### Spindle assembly checkpoint and DNA damage checkpoint are activated upon NPP-3 depletion

Based on Figure 5A and Figure 6A, NPP-3 depletion not only activates the spindle assembly checkpoint (SAC) during metaphase but also induces a redistribution of MDF-1 in prophase. The release of MDF-1 from the NE has been shown to facilitate earlier SAC activation by recruiting MDF-1/MDF-2 to assembled kinetochore prior to nuclear envelope breakdown in human cells (53). In *C. elegans* embryos, similar observations is noted upon of NPP-5 (an cytoplasmic ring Nup), where MDF-1 loses its nuclear envelope (NE) localization in prophase, and accumulates around chromosomes during metaphase, mirroring the effects seen with NPP-3 depletion (52). However, unlike NPP-3 depletion, chromosomes do not localize to nuclear periphery upon NPP-5 depletion (52), and it remains unclear whether the SAC is activated after NPP-5 depletion. Therefore, the loss of MDF-1 localization at the NE is not directly causing peripheral chromosome positioning.

However, with the extension of prophase in NPP-3 depletion dependent on MDF-1 and MDF-2, we suspect that SAC may be activated in prophase, potentially also through the mitotic checkpoint complex (MCC) (72), and this SAC activation could be required for the localization of condensed chromosomes at the nuclear periphery. We hypothesize that early condensation of chromosomes in prophase in NPP-3 depletion may mimic a prometaphase environment, but the delayed outer kinetochore protein assembly could create a microenvironment resembling prometaphase chromosomes with unattached kinetochores as large MDF-1 foci was found to localize around chromosomes in prophase (Figure 5C), potentially resulting in the detection of these chromosomes as “unattached”.

Additionally, NPP-3 depletion activates the DNA damage checkpoint (Figure S3F), likely due to DNA damage caused by compromised nuclear envelope integrity. Since both SAC and DNA damage response components such as MDF-1 and HUS-1 form foci at the nuclear periphery (Figures S3F and 5C), there may be a potential interplay between SAC activation and the DNA damage response following nuclear rupture. For example, MAD1/MAD2 have been implicated in DNA damage responses in both *C. elegans* and human cells during prophase, with RAD-51 foci associated with MDF-2/MAD2 in the proliferative zone of worm germlines (73). However, without MDF-1 in NPP-3 depletion, there are even more HUS-1 foci (Figure S7), suggesting that more DNA remains unrepaired without SAC activation.

The survival of *C. elegans* embryos through anoxia rely on the SAC components SAN-1/MAD-3, MDF-1 and MDF-2 (39). However, whether the prophase chromosome localization to the nuclear periphery is dependent on the SAC has not been analyzed. Investigating the functions of SAC components under various stresses, such as nuclear envelope rupture and DNA damage, beyond the regular detection of unattached kinetochores for prometaphase extension, will be a valuable avenue for future research to explore additional functions of SAC.

## Materials and Methods

### Experimental model and study participant details

*C. elegans* strains

All C. elegans strains were grown at 22°C on Nematode Growth Medium (NGM) plates seeded with *Escherichia coli strain OP50*. All strains used in this study are listed in Table S1.

Transgenic strain construction

The CRISPR-cas9 transgenic strains of endogenous NPP-3::mCherry was generated by SunyBiotech Company [strain name and genotype: PHX913 npp-3(syb913)]. The sequencing primers are listed in Table S2

### Method details

#### RNAi

We performed feeding RNAi experiments using the method previously described (74). The RNAi plasmids with targeted genes were constructed. For single gene knockdown, we inserted DNA fragment of targeted genes into the vector pL4440. For double depletion, the same DNA fragment was inserted into the *npp-3(RNAi)* vector. Primers containing restriction enzyme (RE) for cloning the gene fragment are listed in Table S2.

Clones were verified by single clone PCR of NPP-3 and candidate gene and Sanger Sequencing. The RNAi plasmid was then transformed into HT115 *E. coli* competent cells. The HT115 bacteria with the RNAi plasmid were then seeded on RNAi plates (NGM plates with 100μg/ml ampicillin and 0.1 mM IPTG).

For feeding RNAi experiments, L4 larvae were transferred to NGM plates seeded with corresponding RNAi bacteria for 26 hrs, except for *npp-7(RNAi)*, where late L4 larvae were picked and incubated for 18 hrs. For *baf-1(RNAi)*, we pickedL4 on the RNAi plates and then incubated at 20°C for 48 hrs before microscopy imaging (59).

Embryonic viability and brood size assay

The embryonic viability and brood size assays were performed on the worms fed with HT115 bacteria containing pL4440, and pL4440 inserted NPP-3, MDF-1, or ligated NPP-3 and MDF-1 plasmid to knock down single or double genes. The protocol was adapted from (75). The worms were transferred to fresh plates every 12 hours and the plates were scored. The brood size is defined by the total numbers of embryos and larva on the plates. Embryonic inviability is defined by the number of embryos that did not hatch in 24 hours. There are 3 biological replicates in each experiment, and the experiments were repeated twice.

Sample preparation for RNA Sequencing

Samples for the RNA sequencing are extracted from the embryos (Adapted from Zamanian Lab’s protocol https://www.zamanianlab.org/ZamanianLabDocs/protocols/Caenorhabditis_elegans/RNA_Extra ction/RNA_Extraction/#rna-isolation-from-trizol-samples). The RNA samples were sent to BGI for RNA sequencing and analyses.

Immunofluorescence

N2 fed with *L4440(RNAi)* and *npp-3(RNAi)* bacteria for 26 hrs were prepared. The gravid hermaphrodites were dissected, and embryos were released. The immunofluorescence steps were previously described (76). Primary antibody H3K9me3 (1:500 diluted with antibody dilution buffer, Abcam ab8898) and secondary antibody: goat-anti-Mouse IgG- FITC (1:500, diluted with 1X PBST, Jackson Immuno Research 115-096-062).

Confocal microscopy

Embryos were dissected and mounted in M9 buffer on the 2% agarose pad. The slides were then sealed with Vaseline to prevent evaporation.

For the time-lapse images, images were captured using a Nikon spinning disk confocal microscope equipped with a 100×, 1.4 NA objective lens, and images were acquired with MetaMorph software. To ensure consistency across experiments, parameters such as laser power, exposure time, time interval, and z-stack interval were kept constants for all groups.

The permeability ability experiment (Figure S2) was carried out with DeltaVision (DV) imaging system (60×1.42 NA oil objective).

### Image quantification and Imaris simulation

Quantification of chromosome condensation during cell cycle:

The chromosome condensation assay was based on Maddox’s previous study (41). Z-stack images of GFP::H2B including whole nucleus (12 μm) were acquired from telophase of P0 cell to metaphase of P1 cell. Maximum intensity projection was applied to each frame. As nucleus of P1 cell enlarges during prophase, individual scaling was applied to square region that fit within the nucleus during each time point. This approach ensures that the intensity distribution (0–255) at each timepoint remains independent of the nucleus size or photobleaching effects. The values of pixels in each intensity (0–255) can be acquired by the histogram in ImageJ software and then the total values within various thresholds can be acquired. The condensation parameter under the specific threshold can be calculated as the total values of pixels (below the threshold) / total values of pixel in the square. 35% threshold (gray values in 0-90) was the optimal one to present the difference between the control and *npp-3(RNAi)* groups.

Quantification of nuclear envelope rupture size:

The middle plane of nucleus expressing NPP-1::GFP and H2B::mCherry was selected for analysis. The raw confocal 16-bit images were input to ImageJ. We scaled the intensity to 8-bit, such that the minimum intensity is 0 and maximum intensity is 255 at sites with the highest intensity. By drawing the circle fitting the nuclear envelope, and then using the function (Area to line) to transfer the circle into a line (1-pixel width). The intensity of NPP-1::GFP can be profiled along the line. The gap size was measured by the distance between the neighboring peaks at the value of 255. For the statistics, gap sizes within 2 pixels were discarded. The nuclear gap per nucleus was calculated as the total gap size along the line over the total distance of the line.

Quantification of fluorescence intensity:

For the quantification of MDF-1, MDF-2, HCP-3, KNL-1, HCP-1, and BUB-1 intensities, all images were acquired under identical microscopy parameters to ensure consistency. Z-stack images were processed using maximum intensity projection (58). For analysis, a 40×40 pixel region encompassing the nucleus (containing the MDF-1 signal) was cropped, as well as an adjacent 80×80 pixel region encompassing the nucleus, with surrounded by background.

The mean intensity of background was calculated as: 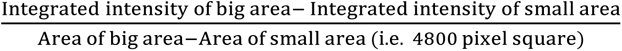

The integrated intensity of the target gene signal was determined by: (Mean intensity of small area- mean intensity of background)*1600 pixel square.

### Statistical analysis

All analyses were done using GraphPad Prism. For comparison between two normally distributed datasets, Student’s t-test was used. For comparison between multiple datasets, one-way ANOVA non-parametric test was used, otherwise two tailed Fisher’s Exact Test was used in the Figure 6D. In all cases, the statistical tests used, the number of samples analyzed, and the p-values are indicated in the figure legends.

## Supplemental information

Document S1. Figures S1–S7 and Table S1-S2

## Acknowledgments

We thank the Yuen lab and Tse lab members for suggestions. We thank the Karen Oegema and Arshad Desai labs for providing strains. We thank the University of Hong Kong School of Biological Sciences Core Facilities, the Li Ka Shing Faculty of Medicine Core Facility, and SUSTech Core Research Facilities for providing equipment for this study. Financial support for this research was provided by General Research Fund (GRF) 17118623 from the Research Grant Council (RGC) of Hong Kong.

## Author contributions

L.J., Y.C.T. and K.W.Y.Y. conceived the study. L.J. performed the methodology and acquired the data. L.J., Y.C.T. and K.W.Y.Y. analyzed and interpreted the data. L.J. wrote the original draft of the manuscript. L.J., Y.C.T. and K.W.Y.Y. reviewed and edited the manuscript. Y.C.T. and K.W.Y.Y. supervised the study.

---

**Figure S1.**
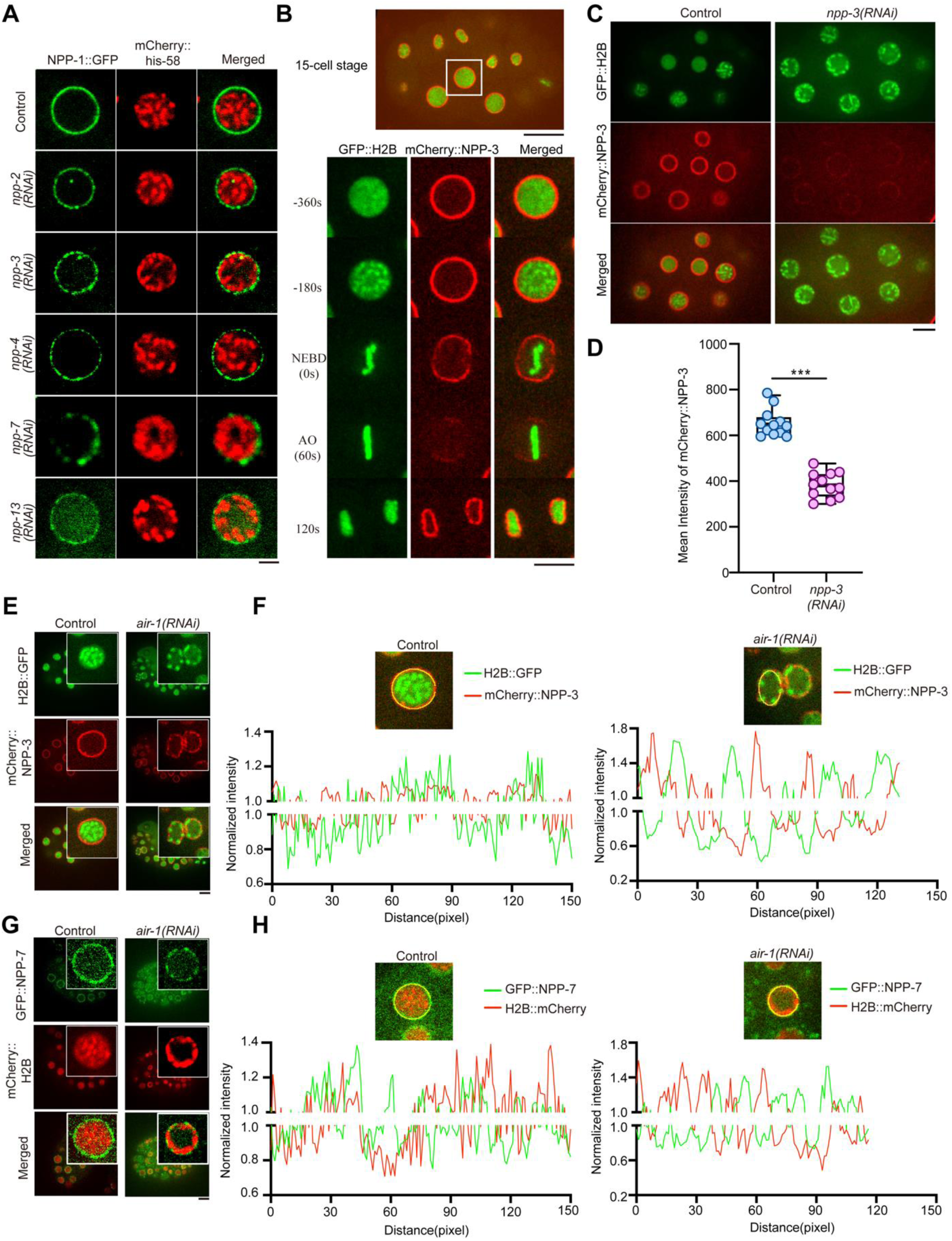
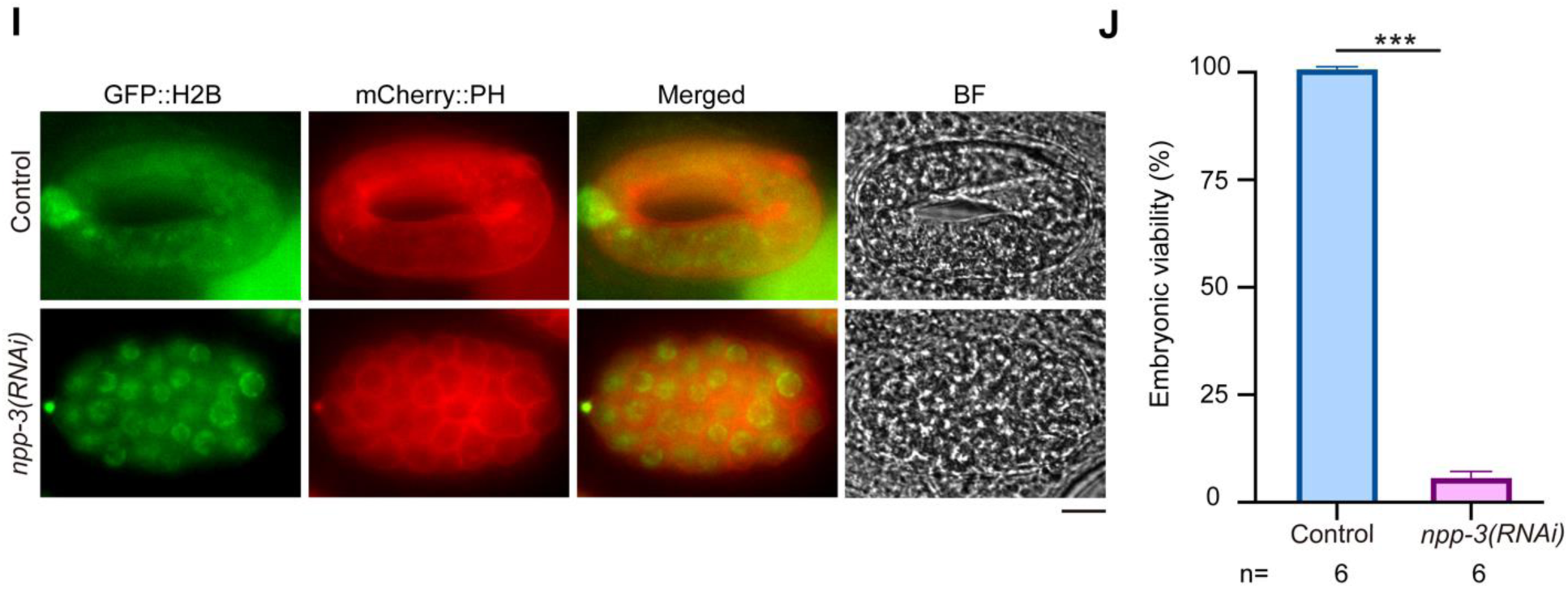
NPP-3 is essential for embryonic development. (A) Representative confocal images in P1 cell of the control, *npp-2(RNAi), npp-3(RNAi), npp-4(RNAi), npp-7(RNAi)*, and *npp-13(RNAi)* embryos expressing NPP-1::GFP and mCherry::H2B. Scale bar 5 μm. (B) Representative confocal image (single layer) of GFP::H2B and mCherry::NPP-3 in the 15-cell stage from prophase to anaphase. White area in the top plane is the time-lapse region for the bottom plane, Scale bar,10 μm for top plane and 5 μm for the bottom one. (C) Representative confocal image (single layer) of GFP::H2B and mCherry::NPP-3 in the 15-cell stage in control and *npp-3(RNAi)* embryos. Scale bar,10 μm. (D) Quantification of mCherry::NPP-3 intensity in the (C). (E) (G) Selected representative confocal images of all chromosomes localizing at the nuclear periphery in the control and *air-1(RNAi)* embryos expressing H2B::GFP and mCherry::NPP-3 (E) and GFP::NPP-7 and mcherry::H2B (G). The upper right image is the zoom-in view of the nucleus. Scale bar 10 μm. (F) (H) The intensity of GFP and mCherry is normalised to the average intensity along the nuclear envelope and plotted in the control and *air-1(RNAi)* nucleus from (E) and (G). (I) Representative confocal image of embryos expressing GFP::H2B and mCherry::PH after 15 hours of imaging from P0 cell in the control and *npp-3(RNAi)* groups. Scale bar 10 μm. (J) Quantification of embryonic viability in the control and *npp-3(RNAi)* embryos.

**Figure S2.**
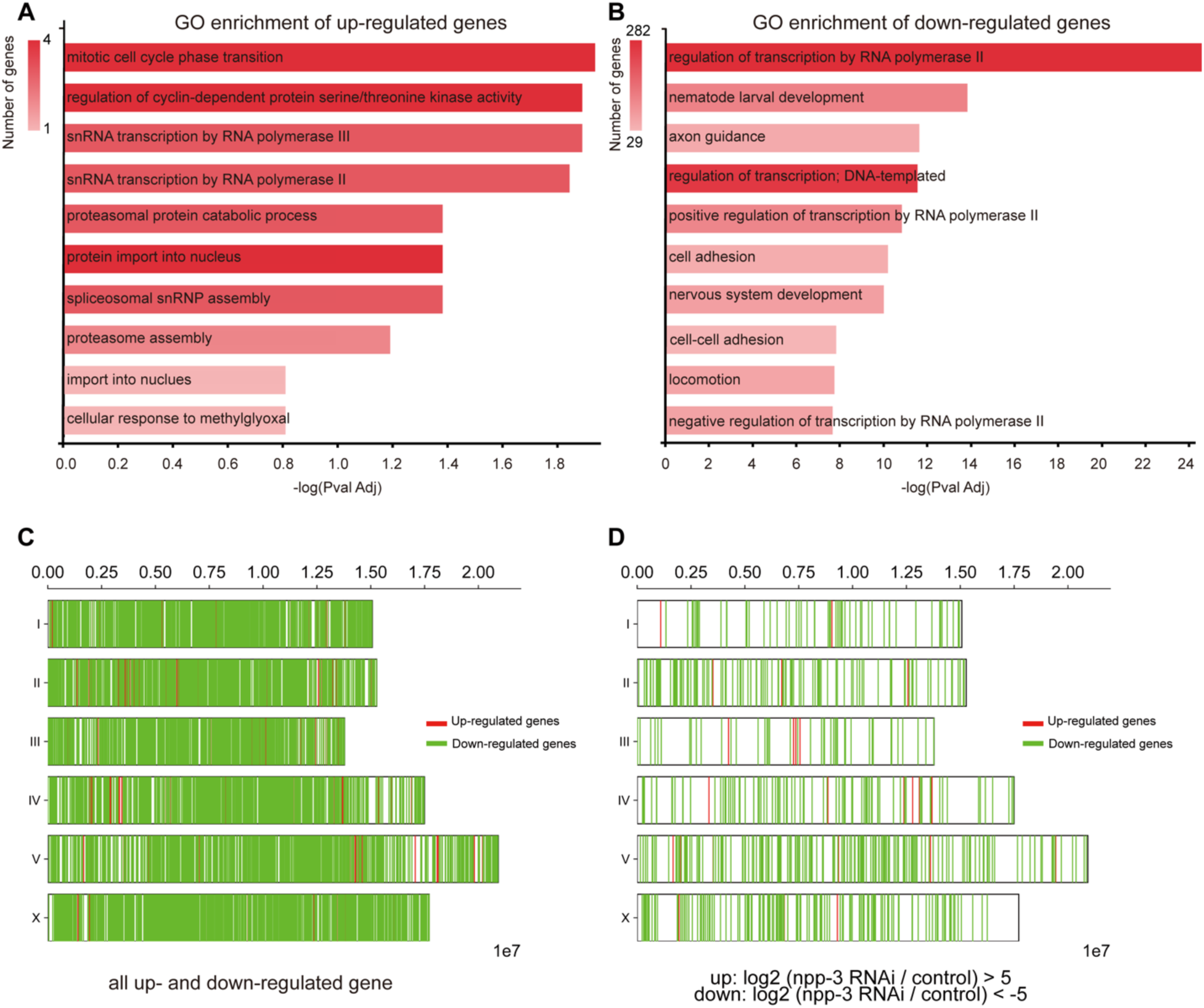
Chromosome localizing to nuclear periphery caused by NPP-3 depletion are transcriptionally repressed. (A) and (B) GO enrichment of up-regulated genes (A) and down-regulated genes (B) in biological process. (C) and (D) Alignment of up-and down regulated genes to chromosome location in different fold- change threshold.

**Figure S3.**
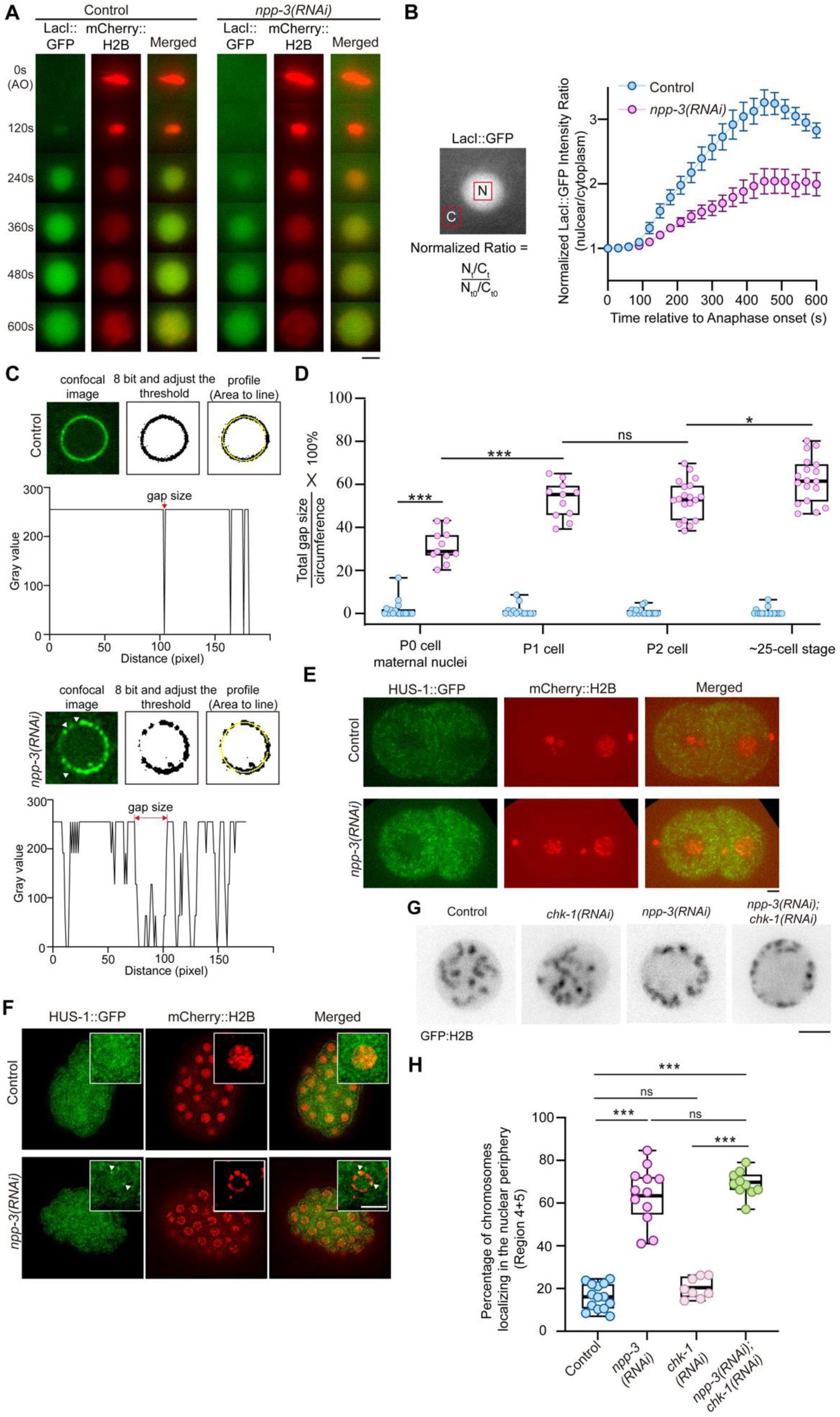
NPP-3 depletion disrupts nuclear envelope integrity and induces DNA damage. (A) Representative maximum projection time-lapse images of LacI::GFP at different time points following anaphase onset in the P1 cell. Scale bar 5 μm. (B) Quantification of LacI intensity during the nuclear envelope reassembly process in the P1 cell. Mean with SEM. (C) Representative single-plane confocal images of P1 cells expressing NPP-1::GFP in control (left panel) and *npp-3(RNAi)* groups (right panel). The raw confocal image was set to 8 bits and subjected to thresholding. This process simplified the intensity values, setting the minimum to 0 and the maximum to 255. The gap size was defined as the distance between the two neighboring peaks at 255. (D) Quantification of total gap size per nucleus in the different cell stages. Each dot represents one nucleus. Error bars represent mean ± SEM. Data were analyzed using Student’s t-test; *p<0.05, **p<0.005, ***p<0.001. (E) Representative maximum projection images of P1 cell expressing HUS-1::GFP and mCherry::H2B in control and *npp-3(RNAi)* groups, Scale bar 5 μm. (F) Representative single-plane images of 20-30 cell stage embryos expressing HUS-1::GFP and mCherry::H2B in control and *npp-3(RNAi)* groups. The upper right image is the zoom-in view of the nucleus. Scale bar 5 μm. (G) Representative single plane images of P1 cell with GFP:::H2B marked chromosomes in control, *npp-3(RNAi),chk-1(RNAi)* and *npp-3(RNAi);chk-1(RNAi).* Scale bar 5 μm. (H) Quantification of the chromosomal nuclear periphery localization in different conditions shown in the (G) by concentric assay shown in Figure 1B.

**Figure S4.**
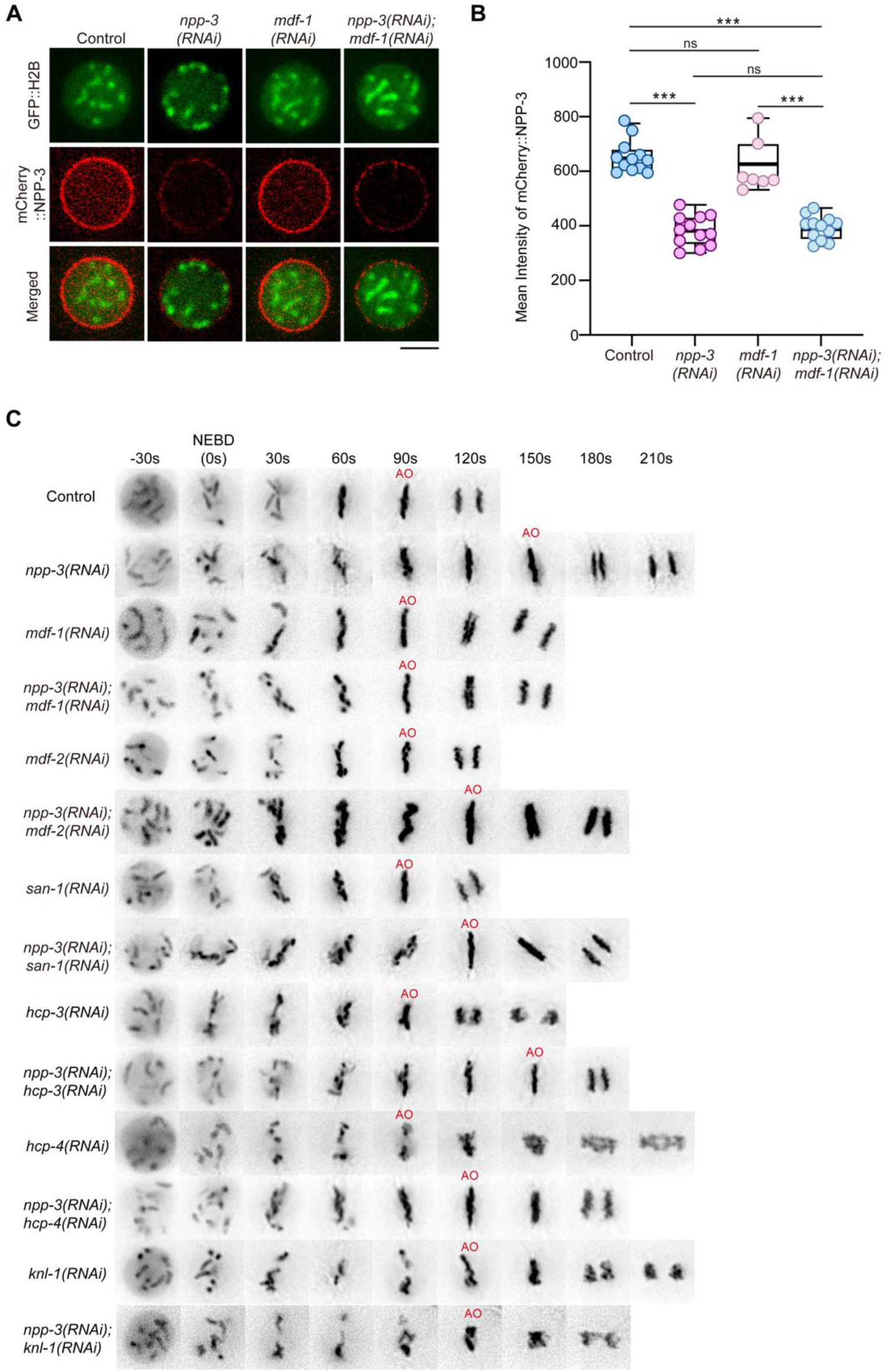
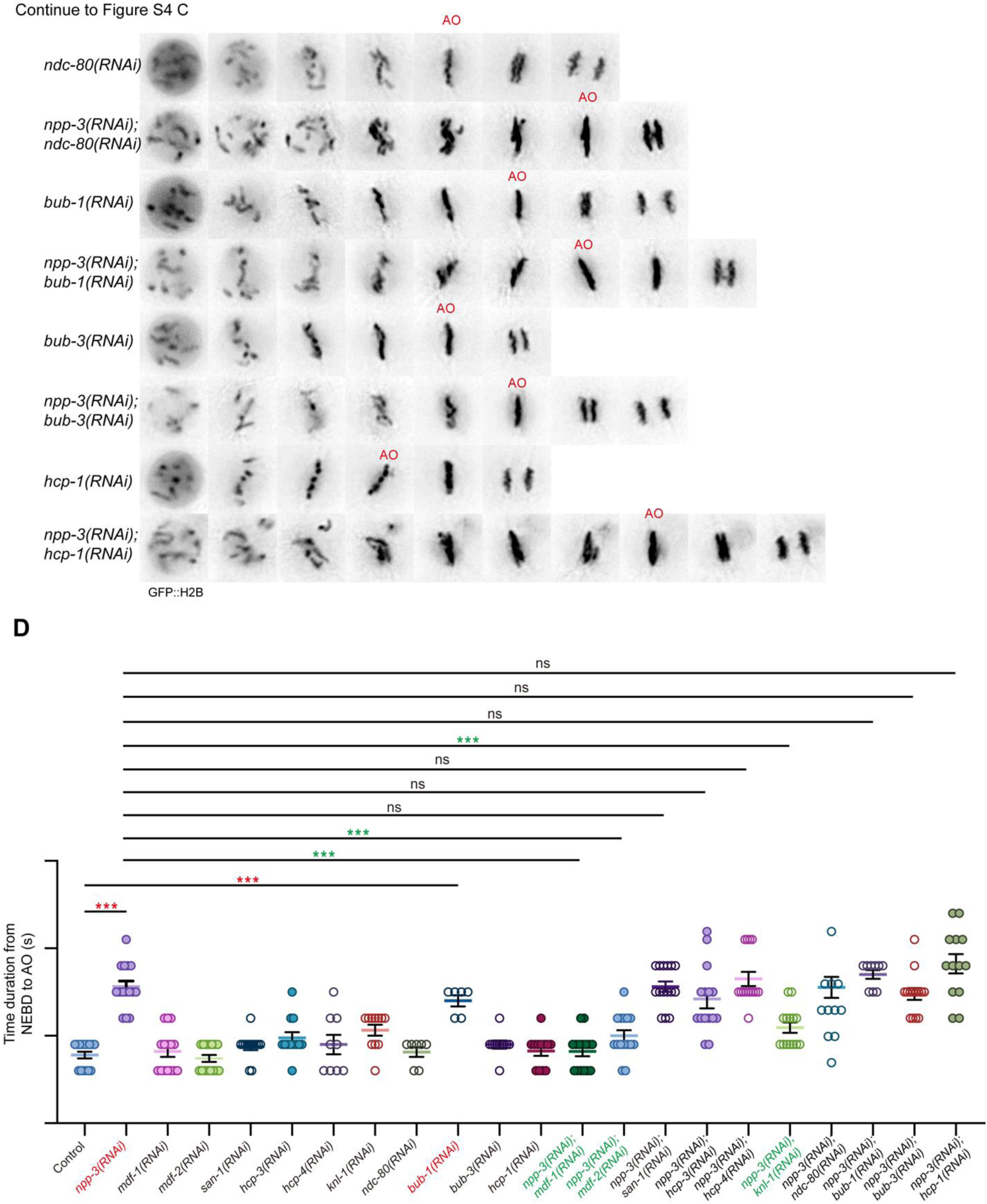
MDF-1, MDF-2 and KNL-1 are required for SAC activation at prometaphase upon NPP-3 depletion. (A) Representative confocal image (single layer) of GFP::H2B and mCherry::NPP-3 in the P1 cell in control, *npp-3(RNAi)*; *mdf-1(RNAi)* and *npp-3(RNAi)*; *mdf-1(RNAi)* groups. Scale bar 5 μm. (B) Quantification of NPP-3 intensity at the nuclear envelope in the (A). Error bars represent mean ± SEM. Data were analyzed using Student’s t-test;***p<0.001, ns: not significant. (C) Representative time-lapse image of GFP::H2B in the P1 cell from prophase to anaphase in different conditions (Figure 3I and 3G). Scale bar 5μm. (D) Quantification of the duration of P1 cell from nuclear envelope breakdown (NEBD) to anaphase onset (AO) in (C). Each dot represents one embryo. Error bars represent mean ± SEM. Data were analyzed by one-way ANOVA (and nonparametric or mixed); *p<0.05, ***p<0.001, ns: not significant. Groups showing a significant difference compared to control are marked in red, while groups with significant differences compared to *npp-3(RNAi)* are marked in green.

**Figure S5.**
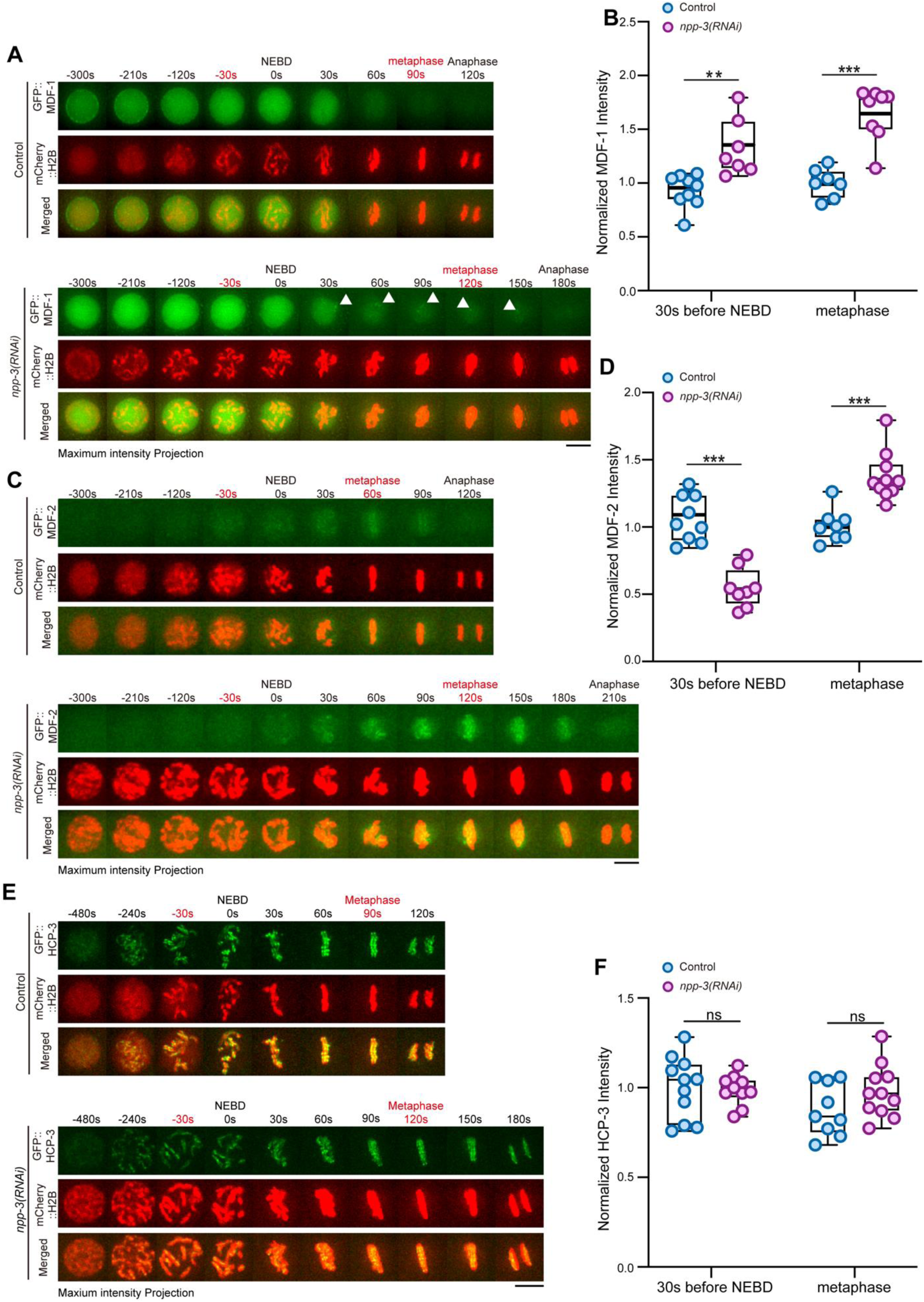
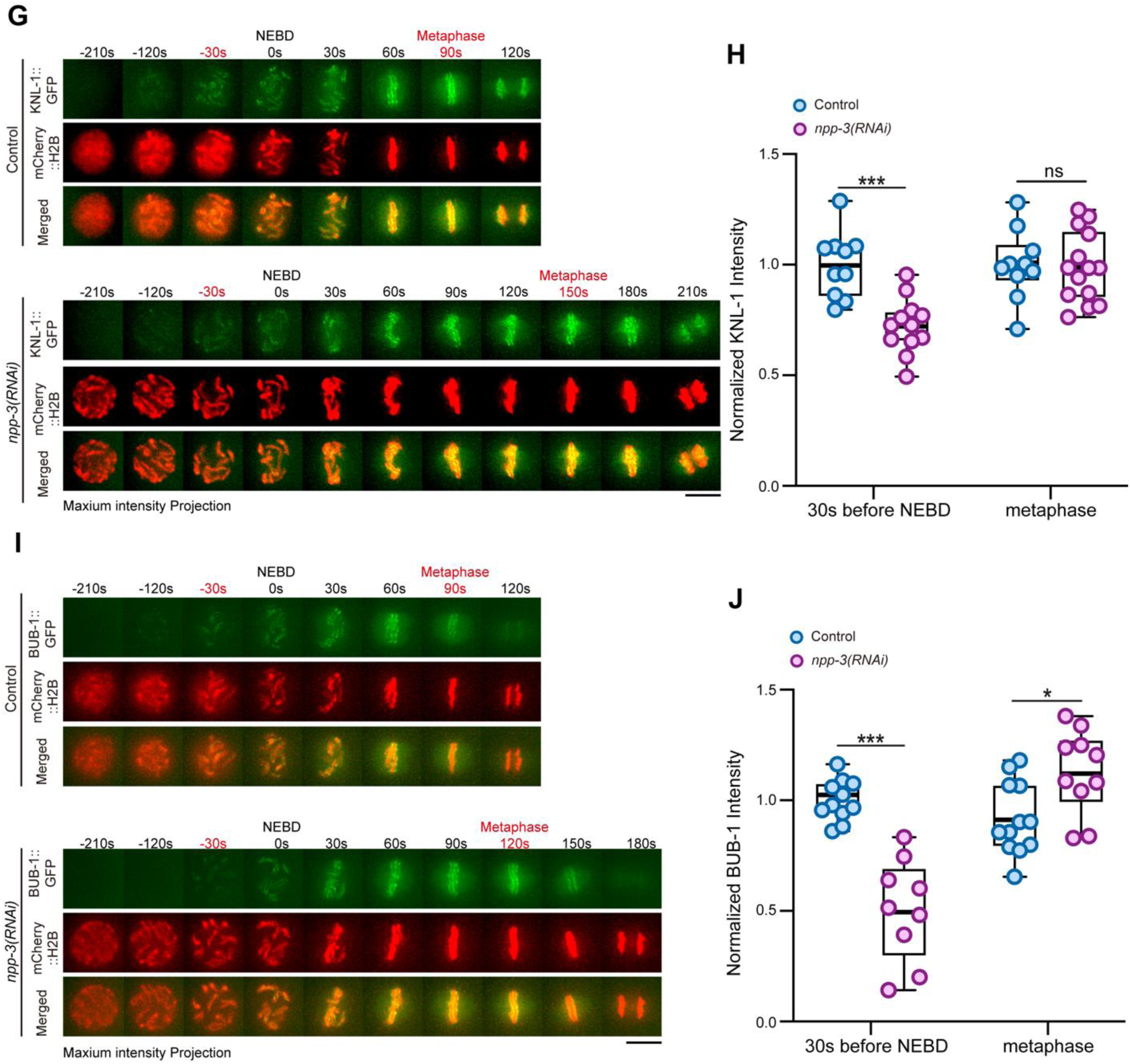
NPP-3 depletion reduces the import of KNL-1, BUB-1 and HCP-1 during prophase in P1 cell. (A) Representative time-lapse image (maximum intensity projection) of GFP::MDF-1 and mCherry::H2B in the EMS cell of control and *npp-3(RNAi)* embryos. Arrows indicate MDF-1 accumulation on chromosomes. Scale bar 5 μm. (B) Quantification of MDF-1 intensity at 30s before NEBD and metaphase in the (A). Error bars represent mean ± SEM. Data were analyzed using student’s t-test, **p<0.005, ***p<0.001. (C) Representative time-lapse image (maximum intensity projection) of GFP::MDF-2 and mCherry::H2B in the EMS cell of control and *npp-3(RNAi)* embryos. Scale bar 5 μm. (D) Quantification of MDF-2 intensity at 30s before NEBD and metaphase in the (C). Error bars represent mean ± SEM. Data were analyzed using student’s t-test, ***p<0.001. (E) (G) (I) (K) Representative time-lapse image (maximum intensity projection) of GFP::SAN-1(E), GFP::HCP-3 (G), KNL-1::GFP (I) and BUB-1::GFP (K) with mCherry::H2B in the P1 of control and *npp-3(RNAi)* embryos. Scale bar 5 μm. (F) (H) (J) (L) Quantification of SAN-1(F), HCP-3 (H), KNL-1 (J) and BUB-1 (L) intensity at 30s before NEBD and metaphase in the (E), (G) (I) (K). Error bars represent mean ± SEM. Data were analyzed using student’s t-test. *p<0.05, **p<0.005, ***p<0.001.

**Figure S6.**
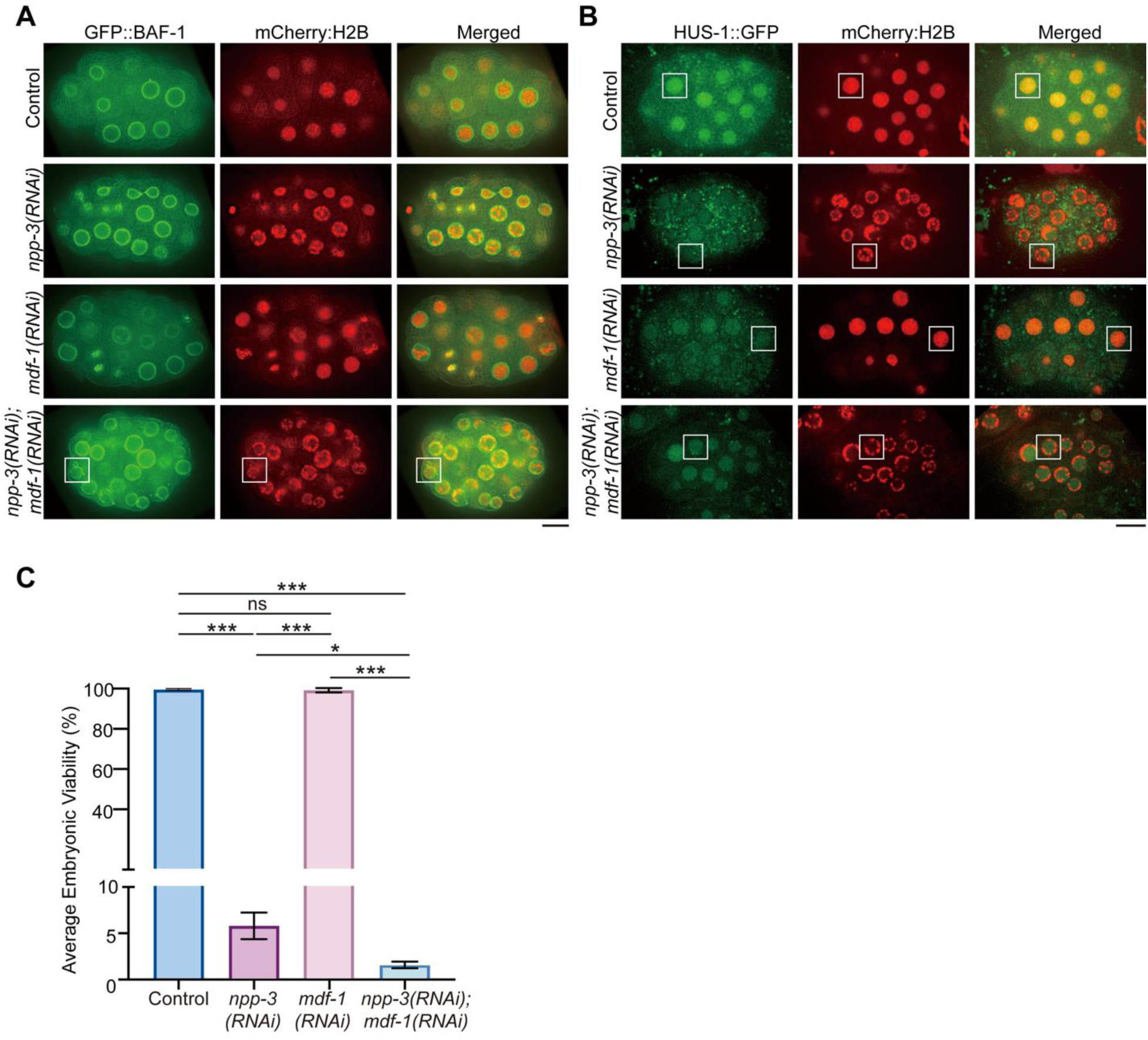
Loss of MDF-1 in NPP-3 depletion leads to more nuclear fragmentation, DNA damage, and higher lethality. (A) Representative single-plane confocal images of 20-30 cell-stage embryos expressing GFP::BAF-1 and mCherry::H2B in control, *mdf-1(RNAi)*, *npp-3(RNAi)*, and *npp-3(RNAi)*; *mdf-1(RNAi)* conditions. White area indicates the micronuclei shown in the Figure 7A. Scale bar 10 μm. (B) Representative images of HUS-1 and chromosome localization in the embryonic stage (about 20-30 cell stage). The white area is the region of magnified confocal images of in the Figure 7B. Scale bar 10 μm. (C) The average embryonic viability in the control, *mdf-1(RNAi)*, *npp-3(RNAi)*, and *npp-3(RNAi)*; *mdf-1(RNAi)* conditions. The data were collected from three plates in each group, with three worms analyzed per plate. Experiments were performed twice.

**Figure S7.**
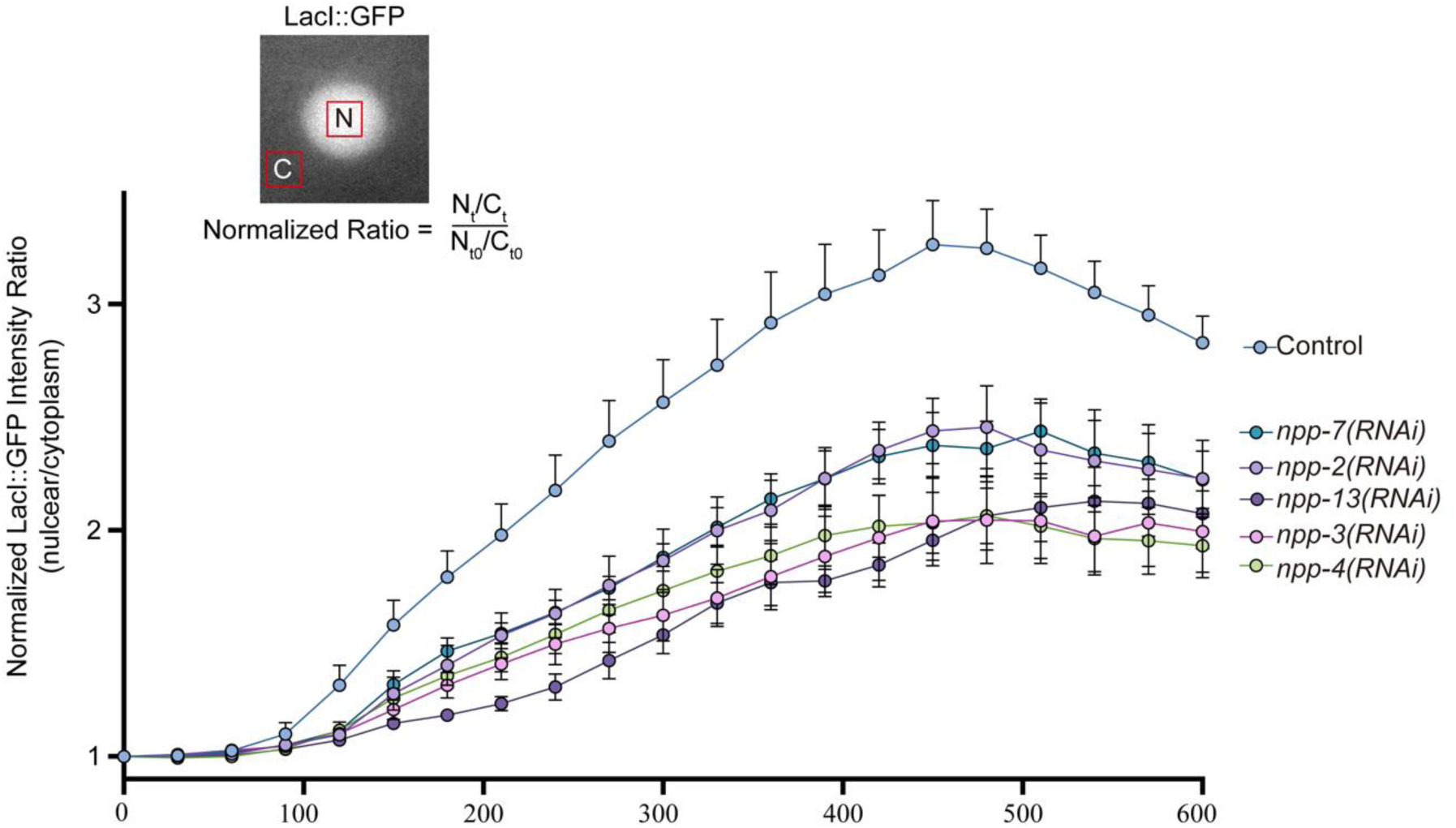
Nuclear peripheral chromosome localization is not solely depending on the nuclear envelope rupture. Normalized intensity ratio LacI::GFP (nuclear versus cytoplasmic) changes over time following anaphase onset in P0 cells of control, *npp-2(RNAi), npp-3(RNAi), npp-4(RNAi), npp-7(RNAi)*, and *npp-13(RNAi)* groups. Error bars show mean ± SEM. Sample size is 6.

**Supplemental Table S1:**
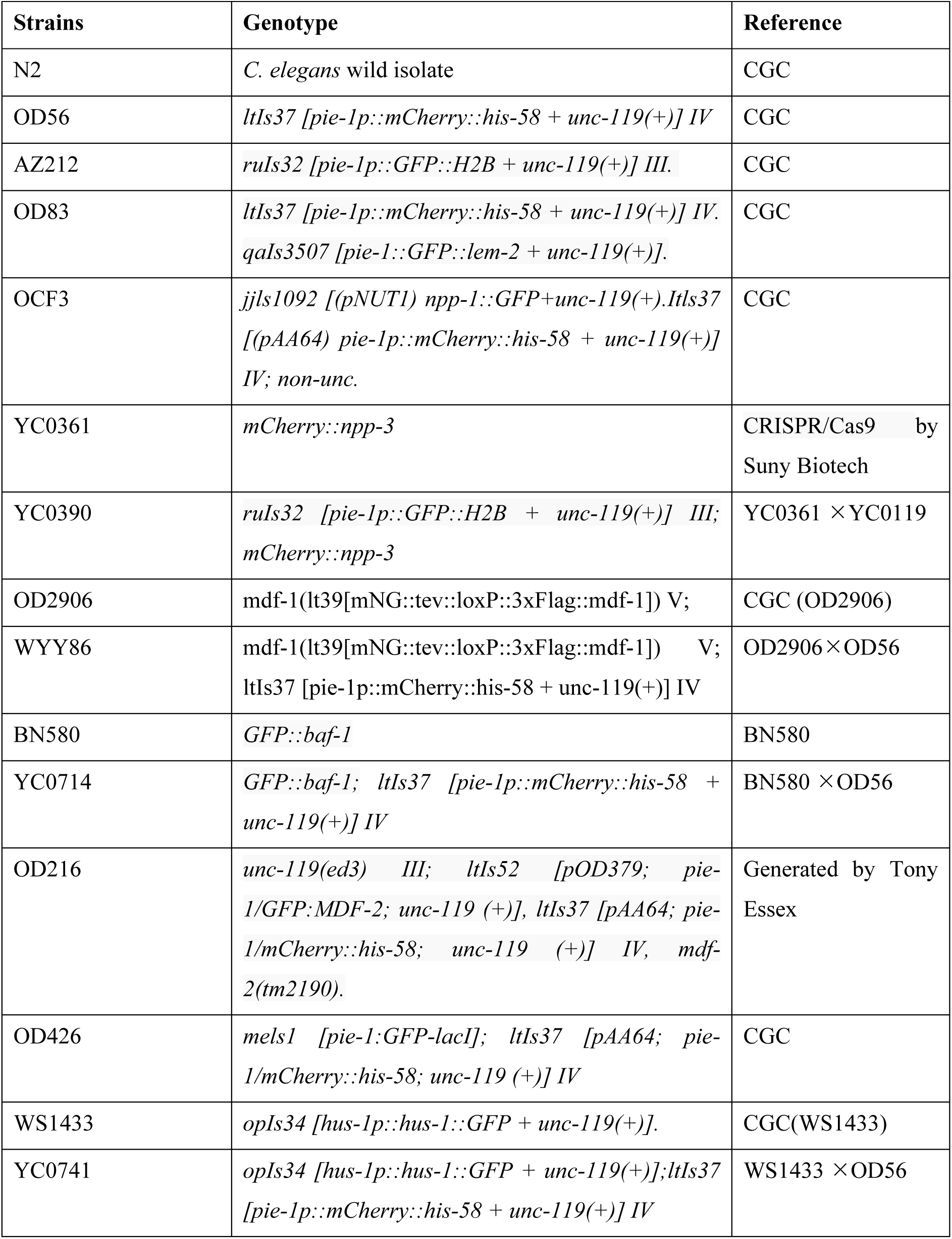
Worm strains.

**Supplemental Table S2:**
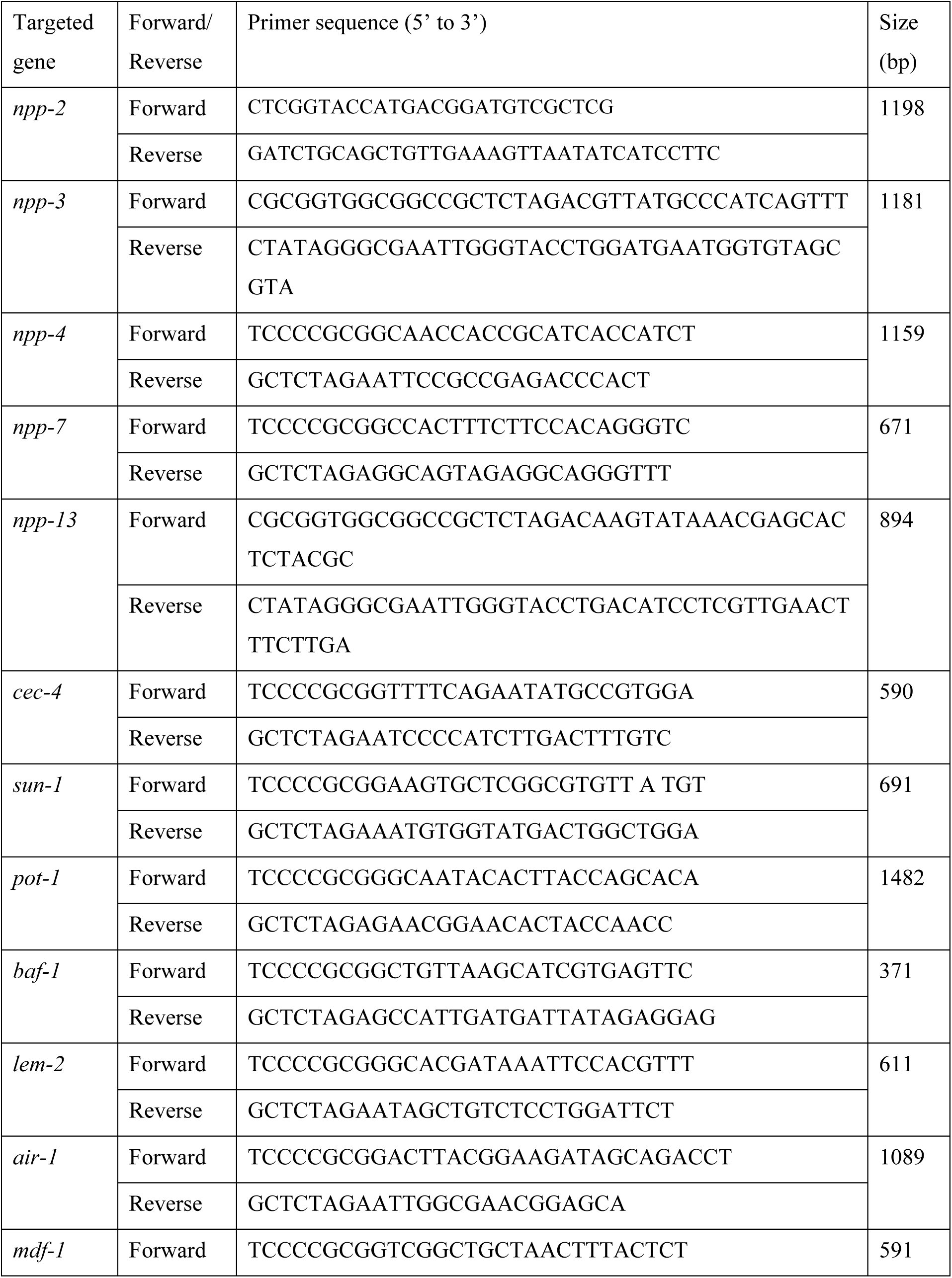
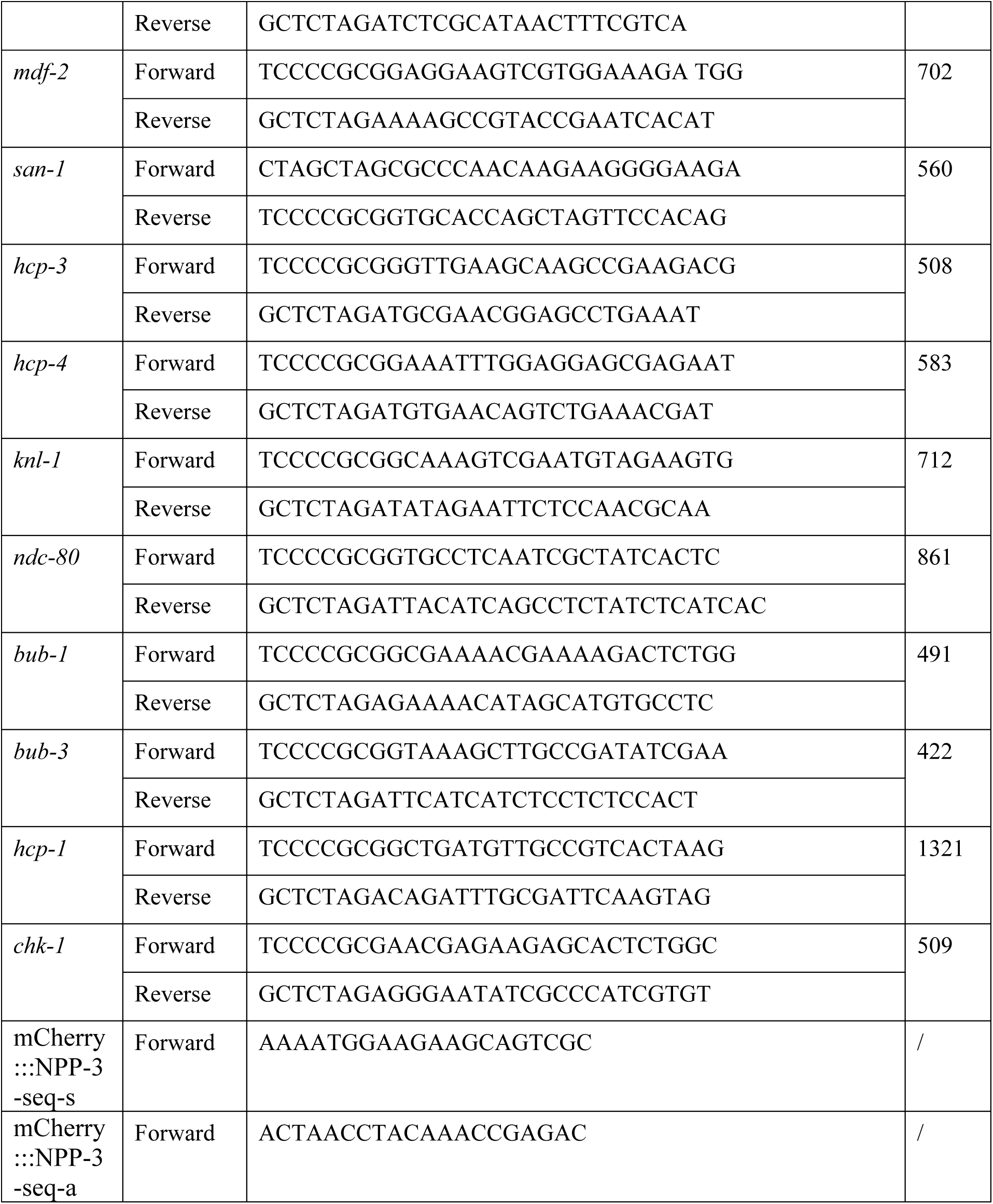
Primer List.

## References

1. Wilson KL, Berk JM. The nuclear envelope at a glance. J Cell Sci. 2010;123(Pt 12):1973–8.

2. Ungricht R, Kutay U. Mechanisms and functions of nuclear envelope remodelling. Nat Rev Mol Cell Biol. 2017;18(4):229–45.

3. Barger SR, Penfield L, Bahmanyar S. Coupling lipid synthesis with nuclear envelope remodeling. Trends Biochem Sci. 2022;47(1):52–65.

4. Burke B, Stewart CL. The nuclear lamins: flexibility in function. Nat Rev Mol Cell Biol. 2013;14(1):13–24.

5. Verstraeten VL, Broers JL, Ramaekers FC, van Steensel MA. The nuclear envelope, a key structure in cellular integrity and gene expression. Curr Med Chem. 2007;14(11):1231–48.

6. Chi YH, Chen ZJ, Jeang KT. The nuclear envelopathies and human diseases. J Biomed Sci. 2009;16:96.

7. Bone CR, Tapley EC, Gorjanacz M, Starr DA. The Caenorhabditis elegans SUN protein UNC-84 interacts with lamin to transfer forces from the cytoplasm to the nucleoskeleton during nuclear migration. Mol Biol Cell. 2014;25(18):2853–65.

8. Li Y, Aksenova V, Tingey M, Yu J, Ma P, Arnaoutov A, et al. Distinct roles of nuclear basket proteins in directing the passage of mRNA through the nuclear pore. Proc Natl Acad Sci U S A. 2021;118(37).

9. de la Cruz Ruiz P, Romero-Bueno R, Askjaer P. Analysis of Nuclear Pore Complexes in Caenorhabditis elegans by Live Imaging and Functional Genomics. Methods Mol Biol. 2022;2502:161–82.

10. Ito N, Sakamoto T, Matsunaga S. Components of the Nuclear Pore Complex are Rising Stars in the Formation of a Subnuclear Platform of Chromatin Organization beyond Their Structural Role as a Nuclear Gate. Cytologia. 2021;86(3):183–7.

11. Grossman E, Medalia O, Zwerger M. Functional architecture of the nuclear pore complex. Annu Rev Biophys. 2012;41:557–84.

12. Richards EJ, Elgin SC. Epigenetic codes for heterochromatin formation and silencing: rounding up the usual suspects. Cell. 2002;108(4):489–500.

13. Babu A, Verma RS. Chromosome structure: euchromatin and heterochromatin. Int Rev Cytol. 1987;108:1–60.

14. Morrison O, Thakur J. Molecular Complexes at Euchromatin, Heterochromatin and Centromeric Chromatin. Int J Mol Sci. 2021;22(13).

15. Parada LA, McQueen PG, Misteli T. Tissue-specific spatial organization of genomes. Genome Biol. 2004;5(7):R44.

16. Shevelyov YY. Interactions of Chromatin with the Nuclear Lamina and Nuclear Pore Complexes. Int J Mol Sci. 2023;24(21).

17. Iglesias N, Paulo JA, Tatarakis A, Wang X, Edwards AL, Bhanu NV, et al. Native Chromatin Proteomics Reveals a Role for Specific Nucleoporins in Heterochromatin Organization and Maintenance. Mol Cell. 2020;77(1):51–66 e8.

18. Dickmanns A, Kehlenbach RH, Fahrenkrog B. Nuclear Pore Complexes and Nucleocytoplasmic Transport: From Structure to Function to Disease. Int Rev Cell Mol Biol. 2015;320:171–233.

19. Vaquerizas JM, Suyama R, Kind J, Miura K, Luscombe NM, Akhtar A. Nuclear pore proteins nup153 and megator define transcriptionally active regions in the Drosophila genome. PLoS Genet. 2010;6(2):e1000846.

20. Labade AS, Karmodiya K, Sengupta K. HOXA repression is mediated by nucleoporin Nup93 assisted by its interactors Nup188 and Nup205. Epigenet Chromatin. 2016;9.

21. Galy V, Mattaj IW, Askjaer P. Caenorhabditis elegans nucleoporins Nup93 and Nup205 determine the limit of nuclear pore complex size exclusion in vivo. Mol Biol Cell. 2003;14(12):5104–15.

22. Hachet V, Busso C, Toya M, Sugimoto A, Askjaer P, Gonczy P. The nucleoporin Nup205/NPP-3 is lost near centrosomes at mitotic onset and can modulate the timing of this process in Caenorhabditis elegans embryos. Molecular Biology of the Cell. 2012;23(16):3111–21.

23. Liu T, Rechtsteiner A, Egelhofer TA, Vielle A, Latorre I, Cheung MS, et al. Broad chromosomal domains of histone modification patterns in C. elegans. Genome Res. 2011;21(2):227–36.

24. Gonzalez-Sandoval A, Towbin BD, Kalck V, Cabianca DS, Gaidatzis D, Hauer MH, et al. Perinuclear Anchoring of H3K9-Methylated Chromatin Stabilizes Induced Cell Fate in C. elegans Embryos. Cell. 2015;163(6):1333–47.

25. Gonzalez-Sandoval A, Gasser SM. Mechanism of chromatin segregation to the nuclear periphery in C. elegans embryos. Worm. 2016;5(3):1–7.

26. Cabianca DS, Munoz-Jimenez C, Kalck V, Gaidatzis D, Padeken J, Seeber A, et al. Active chromatin marks drive spatial sequestration of heterochromatin in C. elegans nuclei. Nature. 2019;569(7758):734–9.

27. Hediger F, Neumann FR, Van Houwe G, Dubrana K, Gasser SM. Live imaging of telomeres: yKu and Sir proteins define redundant telomere-anchoring pathways in yeast. Curr Biol. 2002;12(24):2076–89.

28. Baudrimont A, Penkner A, Woglar A, Machacek T, Wegrostek C, Gloggnitzer J, et al. Leptotene/zygotene chromosome movement via the SUN/KASH protein bridge in Caenorhabditis elegans. PLoS Genet. 2010;6(11):e1001219.

29. Ferreira HC, Towbin BD, Jegou T, Gasser SM. The shelterin protein POT-1 anchors Caenorhabditis elegans telomeres through SUN-1 at the nuclear periphery. J Cell Biol. 2013;203(5):727–35.

30. Crabbe L, Cesare AJ, Kasuboski JM, Fitzpatrick JA, Karlseder J. Human telomeres are tethered to the nuclear envelope during postmitotic nuclear assembly. Cell Rep. 2012;2(6):1521–9.

31. Penkner A, Tang L, Novatchkova M, Ladurner M, Fridkin A, Gruenbaum Y, et al. The nuclear envelope protein Matefin/SUN-1 is required for homologous pairing in C. elegans meiosis. Dev Cell. 2007;12(6):873–85.

32. Hao H, Starr DA. SUN/KASH interactions facilitate force transmission across the nuclear envelope. Nucleus. 2019;10(1):73–80.

33. Halfmann CT, Roux KJ. Barrier-to-autointegration factor: a first responder for repair of nuclear ruptures. Cell Cycle. 2021;20(7):647–60.

34. Maciejowski J, Hatch EM. Nuclear Membrane Rupture and Its Consequences. Annu Rev Cell Dev Biol. 2020;36:85–114.

35. Young AM, Gunn AL, Hatch EM. BAF facilitates interphase nuclear membrane repair through recruitment of nuclear transmembrane proteins. Mol Biol Cell. 2020;31(15):1551–60.

36. Penfield L, Wysolmerski B, Mauro M, Farhadifar R, Martinez MA, Biggs R, et al. Dynein pulling forces counteract lamin-mediated nuclear stability during nuclear envelope repair. Mol Biol Cell. 2018;29(7):852–68.

37. Vinita A. Hajeri BAL, Mary L. Ladage, and Pamela A. Padilla. NPP-16_Nup50 Function and CDK-1 Inactivation Are Associated with Anoxia-induced Prophase Arrest in Caenorhabditis elegans. Molecular Biology of the Cell. 2010;21:712–24.

38. Foe VE. Reversible chromosome condensation induced in Drosophila embryos by anoxia: visualization of interphase nuclear organization. The Journal of Cell Biology. 1985;100(5):1623–36.

39. Nystul TG, Goldmark JP, Padilla PA, Roth MB. Suspended animation in C-elegans requires the spindle checkpoint. Science. 2003;302(5647):1038–41.

40. Schumacher JM, Ashcroft N, Donovan PJ, Golden A. A highly conserved centrosomal kinase, AIR-1, is required for accurate cell cycle progression and segregation of developmental factors in Caenorhabditis elegans embryos. Development. 1998;125(22):4391–402.

41. Maddox PS, Portier N, Desai A, Oegema K. Molecular analysis of mitotic chromosome condensation using a quantitative time-resolved fluorescence microscopy assay. Proc Natl Acad Sci U S A. 2006;103(41):15097–102.

42. Kind J, van Steensel B. Genome–nuclear lamina interactions and gene regulation. Current Opinion in Cell Biology. 2010;22(3):320–5.

43. Harr JC, Gonzalez-Sandoval A, Gasser SM. Histones and histone modifications in perinuclear chromatin anchoring: from yeast to man. EMBO Rep. 2016;17(2):139–55.

44. Skoko D, Li M, Huang Y, Mizuuchi M, Cai M, Bradley CM, et al. Barrier-to-autointegration factor (BAF) condenses DNA by looping. Proc Natl Acad Sci U S A. 2009;106(39):16610–5.

45. Cho S, Vashisth M, Abbas A, Majkut S, Vogel K, Xia Y, et al. Mechanosensing by the Lamina Protects against Nuclear Rupture, DNA Damage, and Cell-Cycle Arrest. Dev Cell. 2019;49(6):920–35 e5.

46. Shankar R, Lettman MM, Whisler W, Frankel EB, Audhya A. The ESCRT machinery directs quality control over inner nuclear membrane architecture. Cell Rep. 2022;38(3):110263.

47. Zheng R, Ghirlando R, Lee MS, Mizuuchi K, Krause M, Craigie R. Barrier-to-autointegration factor (BAF) bridges DNA in a discrete, higher-order nucleoprotein complex. Proc Natl Acad Sci U S A. 2000;97(16):8997–9002.

48. Montes de Oca R, Lee KK, Wilson KL. Binding of barrier to autointegration factor (BAF) to histone H3 and selected linker histones including H1.1. J Biol Chem. 2005;280(51):42252–62.

49. Kwon M, Leibowitz ML, Lee JH. Small but mighty: the causes and consequences of micronucleus rupture. Exp Mol Med. 2020;52(11):1777–86.

50. Gomes ER, de Almeida SF. TREX1-dependent DNA damage links nuclear rupture to tumor cell invasion. Dev Cell. 2021;56(22):3040–1.

51. Lussi YC, Shumaker DK, Shimi T, Fahrenkrog B. The nucleoporin Nup153 affects spindle checkpoint activity due to an association with Mad1. Nucleus. 2010;1(1):71–84.

52. Rodenas E, Gonzalez-Aguilera C, Ayuso C, Askjaer P. Dissection of the NUP107 nuclear pore subcomplex reveals a novel interaction with spindle assembly checkpoint protein MAD1 in Caenorhabditis elegans. Mol Biol Cell. 2012;23(5):930–44.

53. Pines J, Choudhary JS, Tyson AL, Yu L, Pardo M, Barbiero M, et al. Cyclin B1-Cdk1 facilitates MAD1 release from the nuclear pore to ensure a robust spindle checkpoint. Journal of Cell Biology. 2020;219(6).

54. Rodriguez-Bravo V, Maciejowski J, Corona J, Buch HK, Collin P, Kanemaki MT, et al. Nuclear pores protect genome integrity by assembling a premitotic and Mad1-dependent anaphase inhibitor. Cell. 2014;156(5):1017–31.

55. Hajeri VA, Trejo J, Padilla PA. Characterization of sub-nuclear changes in Caenorhabditis elegans embryos exposed to brief, intermediate and long-term anoxia to analyze anoxia-induced cell cycle arrest. BMC Cell Biol. 2005;6:47–61.

56. Pandey R, Heeger S, Lehner CF. Rapid effects of acute anoxia on spindle kinetochore interactions activate the mitotic spindle checkpoint. J Cell Sci. 2007;120(Pt 16):2807–18.

57. Dhanya K. Cheerambathur RG, Brian Cook, Karen Oegema, Arshad Desai. Crosstalk Between Microtubule Attachment Complexes Ensures Accurate Chromosome Segregation. Science. 2013;342 (6163):1239–42.

58. Anthony Essex AD, Lindsay Lewellyn,, Karen Oegema aAD. Systematic Analysis in Caenorhabditis elegans Reveals that the Spindle Checkpoint Is Composed of Two Largely Independent Branches. Molecular Biology of the Cell. 2009;20:1252–67.

59. Gorjanacz M, Klerkx EP, Galy V, Santarella R, Lopez-Iglesias C, Askjaer P, et al. Caenorhabditis elegans BAF-1 and its kinase VRK-1 participate directly in post-mitotic nuclear envelope assembly. EMBO J. 2007;26(1):132–43.

60. Margalit A, Neufeld E, Feinstein N, Wilson KL, Podbilewicz B, Gruenbaum Y. Barrier to autointegration factor blocks premature cell fusion and maintains adult muscle integrity in C. elegans. J Cell Biol. 2007;178(4):661–73.

61. Foley EA, Kapoor TM. Microtubule attachment and spindle assembly checkpoint signalling at the kinetochore. Nat Rev Mol Cell Biol. 2013;14(1):25–37.

62. Moyle MW, Kim T, Hattersley N, Espeut J, Cheerambathur DK, Oegema K, et al. A Bub1-Mad1 interaction targets the Mad1-Mad2 complex to unattached kinetochores to initiate the spindle checkpoint. J Cell Biol. 2014;204(5):647–57.

63. Stear JH, Roth MB. The Caenorhabditis elegans kinetochore reorganizes at prometaphase and in response to checkpoint stimuli. Molecular Biology of the Cell. 2004;15(11):5187–96.

64. Dammermann A, Maddox PS, Desai A, Oegema K. SAS-4 is recruited to a dynamic structure in newly forming centrioles that is stabilized by the gamma-tubulin-mediated addition of centriolar microtubules. J Cell Biol. 2008;180(4):771–85.

65. Gerhold AR, Poupart V, Labbe JC, Maddox PS. Spindle assembly checkpoint strength is linked to cell fate in the Caenorhabditis elegans embryo. Mol Biol Cell. 2018;29(12):1435–48.

66. Pintard L, Bowerman B. Mitotic Cell Division in Caenorhabditis elegans. Genetics. 2019;211(1):35–73.

67. Hofmann ER, Milstein S, Boulton SJ, Ye MJ, Hofmann JJ, Stergiou L, et al. Caenorhabditis elegans HUS-1 is a DNA damage checkpoint protein required for genome stability and EGL-1-mediated apoptosis. Current Biology. 2002;12(22):1908–18.

68. Torfeh E, Simon M, Muggiolu G, Deves G, Vianna F, Bourret S, et al. Monte-Carlo dosimetry and real-time imaging of targeted irradiation consequences in 2-cell stage Caenorhabditis elegans embryo. Sci Rep. 2019;9(1):10568.

69. Gauthier BR, Comaills V. Nuclear Envelope Integrity in Health and Disease: Consequences on Genome Instability and Inflammation. Int J Mol Sci. 2021;22(14).

70. Charar C, Metsuyanim-Cohen S, Gruenbaum Y, Bar DZ. Exploring the nuclear lamina in health and pathology using C. elegans. Curr Top Dev Biol. 2021;144:91–110.

71. Torras-Llort M, Medina-Giro S, Escudero-Ferruz P, Lipinszki Z, Moreno-Moreno O, Karman Z, et al. A fraction of barrier-to-autointegration factor (BAF) associates with centromeres and controls mitosis progression. Commun Biol. 2020;3(1):454.

72. McAinsh AD, Kops G. Principles and dynamics of spindle assembly checkpoint signalling. Nat Rev Mol Cell Biol. 2023;24(8):543–59.

73. Lawrence KS, Chau T, Engebrecht J. DNA damage response and spindle assembly checkpoint function throughout the cell cycle to ensure genomic integrity. PLoS Genet. 2015;11(4):e1005150.

74. Conte D, Jr., MacNeil LT, Walhout AJM, Mello CC. RNA Interference in Caenorhabditis elegans. Curr Protoc Mol Biol. 2015;109:26 3 1- 3 30.

75. Kwah JK, Jaramillo-Lambert A. Measuring Embryonic Viability and Brood Size in Caenorhabditis elegans. J Vis Exp. 2023(192).

76. Wong CYY, Tsui HN, Wang Y, Yuen KWY. Argonaute protein CSR-1 restricts localization of holocentromere protein HCP-3, the C. elegans CENP-A homolog. J Cell Sci. 2024;137(18).

